# From human joints to bioreactor setups: quantifying mechanical stimuli in cartilage physiology and regeneration

**DOI:** 10.1101/2025.10.06.678667

**Authors:** Satanik Mukherjee, Wouter Wilson, Liesbet Geris

## Abstract

**Objective:** Bioreactors are widely used to apply mechanical stimuli to osteochondral (OC) explants and cartilage tissue-engineered (TE) constructs, yet their ability to replicate native joint mechanics is not well quantified. This study benchmarks common bioreactor loading protocols against a finite element (FE) model of the human knee during gait, enabling direct comparison to physiologically relevant mechanical parameters.

**Methods:** A validated FE model of the human knee joint simulating the stance phase of gait was used to characterize physiological mechanical conditions, including maximum principal stress, maximum shear strain, pore pressure, and fluid velocity. These outputs were compared with those generated by representative bioreactor setups: dynamic unconfined compression (10–30%) and combined compression (10%) with ball rotation (±25°), which were applied to both OC plugs and TE constructs, and hydrostatic pressure (0.5–50 MPa), which was applied only to TE constructs. FE simulations evaluated spatial and magnitude-based agreement with native cartilage mechanics.

**Results:** In OC plugs, 10% unconfined compression generated maximum principal stresses (∼7.5 MPa) and pore pressures (∼ 4 MPa) closely matching native tissue (∼ 4.5 MPa and ∼5 MPa, respectively). In TE constructs, even at 30% unconfined strain, maximum principal stresses and pore pressures remained around 100 times lower than physiological values, while fluid velocities were 10 times higher. Hydrostatic loading of TE constructs at 5 MPa matched native pore pressures (∼5 MPa) but induced negligible strains.

**Conclusions:** This study provides a quantitative framework for evaluating how well bioreactor loading regimens replicate physiological joint mechanics. The findings offer actionable guidance to experimentalists for selecting bioreactor conditions based on specific mechanical targets. This can lead to more effective design of *in vitro* cartilage studies and support translational strategies in cartilage repair and tissue engineering.

## 1. Introduction

Mechanical loading plays a crucial role in maintaining articular cartilage homeostasis. Non-physiological mechanical loading, including both hypo-physiological loading, as seen during joint immobilization, and hyper-physiological loading, as encountered during traumatic joint injuries, are linked to degenerative changes in cartilage [1, 2, 3]. Articular cartilage is a biphasic tissue, consisting of a solid phase mainly composed of proteoglycans and collagen fibrils, and a fluid phase. In healthy cartilage, the majority of load-bearing is achieved through pore fluid pressurization. Additionally, there is an arcade-like pattern of collagen fibril orientation across the thickness of cartilage, especially in weight-bearing joints such as the hip and knee. This unique arrangement gives cartilage an anisotropic behavior, allowing it to resist shear forces along the surface. During daily activities, articular cartilage is subjected to a wide range of mechanical stimuli, creating a complex dynamic state of stresses and strains at various anatomical locations within the joint.

In cartilage tissue engineering (TE), bioreactors are used to culture tissue engineered constructs under controlled mechanical conditions. Bioreactors furthermore provide an *ex vivo* platform for studying the effects of mechanical loading on disease progression, such as osteoarthritis (OA) and possible therapeutic interventions. In literature, bioreactors used in context of cartilage (patho)physiology and regeneration can be broadly classified into dynamic compression, combined dynamic compression and shear, and hydrostatic pressure [4, 5]. Each of these bioreactors is designed to mimic a certain aspect of the mechanics encountered *in vivo*. Despite the widespread use of these bioreactors in the literature, there is considerable variability in the corresponding mechanical loading protocols applied in these bioreactors, which can lead to inconsistencies in the outcomes of different studies. While multiple review articles exist that compare these different bioreactors (and loading conditions) to their respective outcomes in terms of cellular response [4, 5], no such broad efforts have been made to compare quantitatively what are the differences in mechanics encountered in these different setups and how closely are they related to the *in vivo* physiological scenario.

Capturing the detailed mechanics of TE constructs or explants subjected to mechanical loading in bioreactors is very difficult using traditional experimental techniques due to the complexity of mechanical environment that exists within the specimens. In this context, *in silico* models have been established as an enabling technology to quantify the mechanical environment in these setups as well as human joints [6, 7]. In particular finite element (FE) analysis has been widely used in literature to quantify the mechanics of constructs or explants in many bioreactor studies [8, 9, 10, 11, 12, 13]. However, most of these studies focus on a single type of bioreactor only, thereby making it difficult to make a comparative analysis across different types of bioreactors and corresponding loading protocols. Furthermore, these studies do not incorporate a model of the *in vivo* scenario which the bioreactor aims to recapitulate as a quantitative basis of comparison.

The present study aims to bridge this gap in literature by performing a comparative analysis of widely used bioreactor setups in the literature with a physiological human knee joint model, utilizing FE approaches. This study will not only provide a better understanding of the differences in mechanical environment across different bioreactor setups, but will also provide an extensive repository of bioreactor setups and loading conditions for experimentalists to choose from, depending on the research question to be studied. To achieve this aim, firstly, a FE element model of the human knee joint is developed using geometry and material properties obtained from literature. Secondly, FE models of bioreactors applying mechanical loading to osteochondral grafts and TE constructs are developed. Lastly, a comparative analysis is conducted between the optimal loading cases of each bioreactor type and the human knee joint. This analysis will help identify the loading regimes specific to each bioreactor setup that most closely mimics specific mechanical aspects of the physiological situation.

## 2. Methods

### 2.1. FE modeling of the human knee joint

The tibiofemoral joint geometry of a cadaveric right knee (70-year-old female, 77.1 kg, 1.68 m) was obtained from the Open Knee database [14] and imported into Abaqus 2023 (SIMULIA, Providence, RI, USA) for FE analysis. Model development details are provided in Supplementary Data S1. Briefly, articular cartilage was modeled as a biphasic, fibril-reinforced poroviscoelastic (FRPVE) material, as described previously [15, 16]. The solid matrix was composed of two main components: collagen fibrils (fibrillar) and proteoglycans (non-fibrillar). A depth-dependent, arcade-like orientation was applied to the primary collagen fibrils to represent their natural organization through the cartilage thickness. Additionally, split-line patterns were incorporated on the cartilage surface to reflect the preferred in-plane orientation of the collagen network (Figure 1(c)). Material parameters for all tissues are listed in Supplementary Table S1.

**Figure 1:**
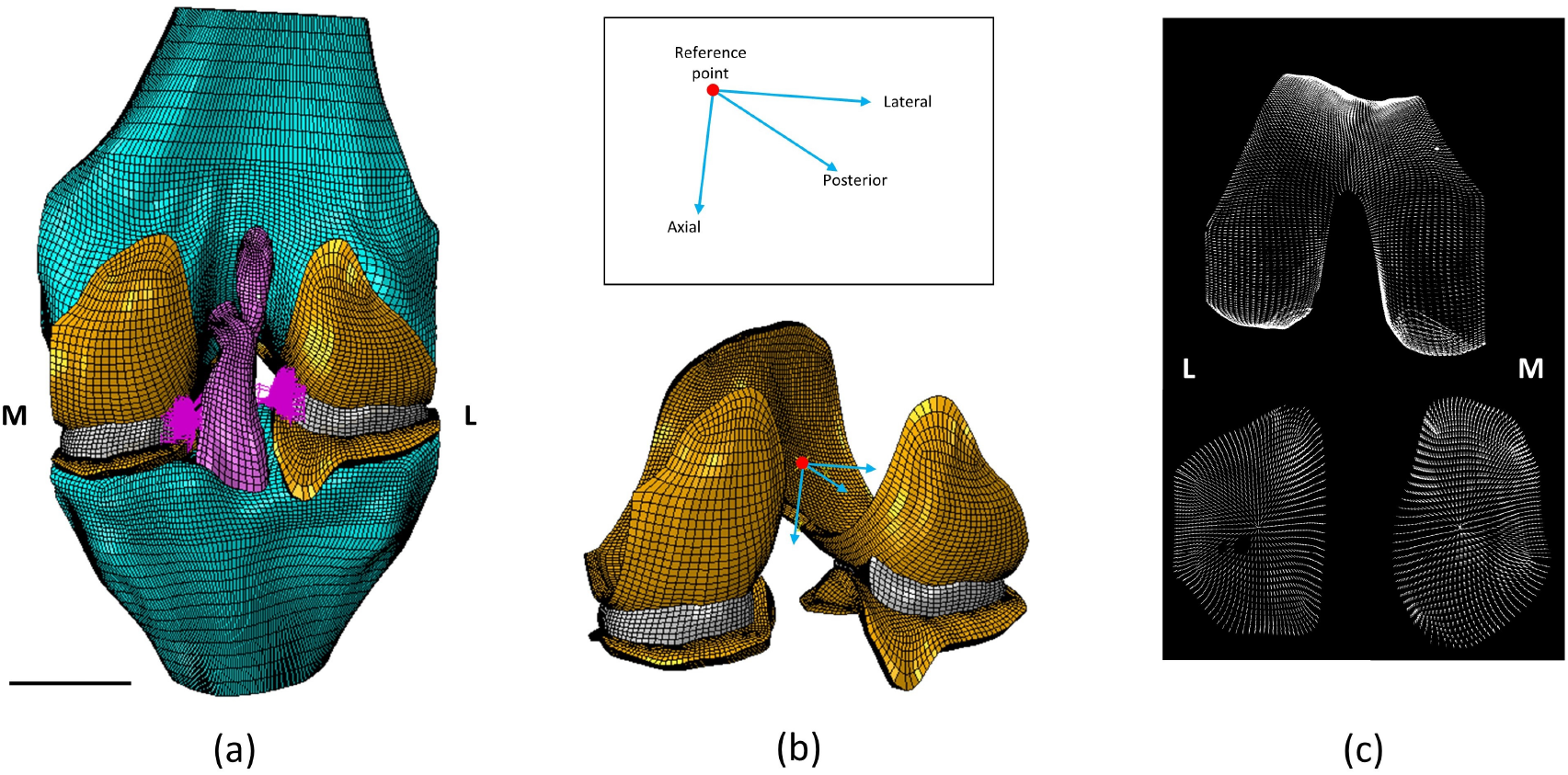
(a) FE model of the human knee joint with the different tissues and meniscal attachment springs (in pink), M-medial, L-lateral, Scale bar = 20 mm; (b) The reference point and the axes about which boundary conditions are applied for the FE model (ligaments and bones have been removed for visualization). In the inset box, a zoomed in view showing the reference point and axes with anatomical directions; (c) Orientation of the split-line patterns at the superficial zone as implemented in the FE model.

Boundary conditions simulating the stance phase of gait, including time-dependent axial force, anterior-posterior translation, and internal–external and flexion–extension rotations—were applied to the femur, based on Mononen et al. [17]. Axial force was scaled by 0.77 to reflect the donor’s body weight. Boundary conditions were applied at a reference point centered between the femoral epicondyles (Figure 1(b)). The tibia was fully constrained.

### 2.2. FE modeling of bioreactors for osteochondral plugs

Three-dimensional FE models of the osteochondral plugs were created in Abaqus, with an 8 mm diameter based on literature [18, 19]. Model development details are provided in Supplementary Data S2. Briefly, the osteochondral plugs consisted of cartilage (2 mm thick considering average human cartilage thickness), calcified cartilage (0.2 mm thick [20]) and 4 mm of subchondral bone underneath. The subchondral bone was further subdivided into subchondral cortical bone (0.36 mm thick) and subchondral trabecular bone (3.64 mm thick) [21]). The corresponding tissues were rigidly attached to one another by using a ‘Tie constraint’. Cartilage was modeled as a fibril-reinforced poroviscoelastic material as described in the previous section. Calcified cartilage, subchondral cortical and trabecular bone were modeled as linear elastic materials with material parameters obtained from literature [22], and mentioned in Supplementary Table S1.

The osteochondral plugs were subject to mechanical loading in two different bioreactor setups that are commonly used in literature to apply unconfined compression and combined compression and shear respectively. The boundary conditions that were applied to mimic mechanical loading in FE model of the bioreactors are described below (Figure 2) :

**Figure 2:**
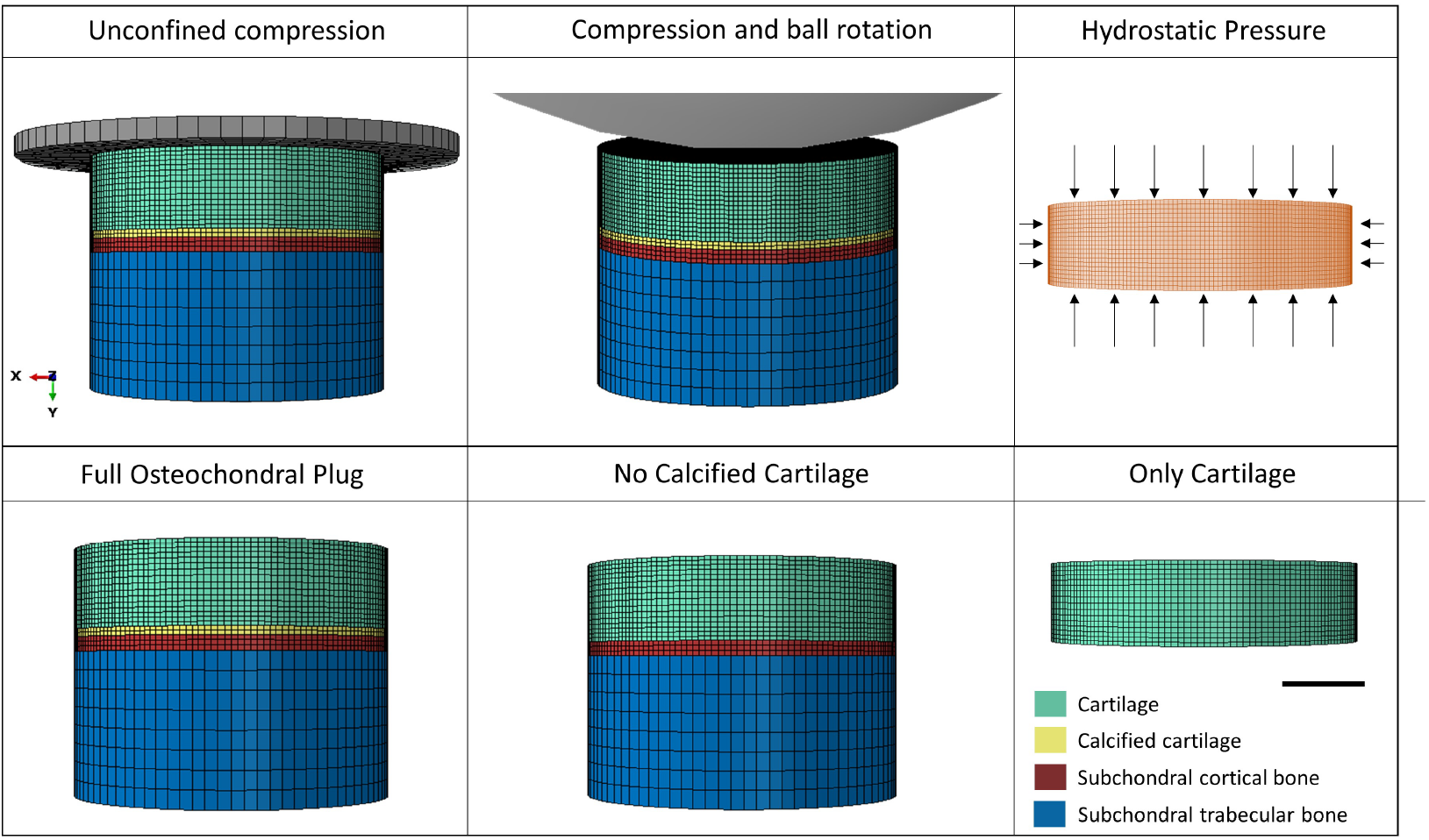
Top row: Different types of bioreactors used in the study. The first two from left show osteochondral plugs, while hydrostatic pressure was applied to only TE constructs. A portion of the ball is visualized in the compression and ball rotation case. Bottom row: FE models to study the role of different tissues in osteochondral plugs. Scale bar = 2 mm.

#### (a) Dynamic unconfined uniaxial compression bioreactor

This bioreactor is most commonly used in literature for its simplicity in design and ease of application of mechanical loading [23, 24]. Uniaxial dynamic compression was applied to the plugs in the FE model by using a rigid impermeable plate with a diameter of 12 mm. A haversine function with a frequency of 1 Hz and amplitudes of 10%, 20%, and 30% of the initial cartilage thickness was defined at the reference point of the rigid plate to impart dynamic compression to the plugs. The loading amplitudes were selected to encompass the range of hypo to hyper-physiological loading regimes mentioned in literature [23, 25].

To assess the role of individual tissues in the stress response of osteochondral plugs subject to uniaxial compression, three cases were simulated by progressively excluding specific tissues (Figure 2):

a. All: included cartilage, calcified cartilage and subchondral bone
b. No calcified cartilage (NoCC): calcified cartilage was excluded
c. Only cartilage: 2 mm thick cartilage layer only

#### (b) Dynamic multi-axial loading bioreactor

To replicate physiological multi-axial loading, bioreactors combining axial compression and shear have been developed in literature, notably using a 32 mm ceramic hip ball as described in studies from AO Davos, Switzerland [26, 27]. In this configuration, axial load is applied via vertical displacement of the ball, while shear is generated by rotating the ball about an axis perpendicular to the loading direction. An FE model of this setup was developed in this study, with the ceramic ball modeled as a rigid body. Boundary conditions were applied to a reference point at the center of the ball, with four loading conditions tested in this setup:

1. 10% quasi-static compression for 10 seconds,
2. 10% quasi-static compression, followed by 10% dynamic loading at 1 Hz,
3. 10% quasi-static compression, followed by ±25 degree dynamic rotation at 1 Hz,
4. 10% quasi-static compression, followed by 10% dynamic loading and ±25 degree dynamic rotation at 1 Hz.

All dynamic conditions were simulated for five cycles in the FE model. Results were analyzed at the peak of the fifth cycle to ensure that the initial transient effects had dissipated.

### 2.3. FE modeling of bioreactors for cartilage tissue engineered constructs

The tissue-engineered (TE) constructs were modeled as 8 mm diameter, 2 mm thick discs. The dimensions were kept consistent with those of the cartilage in osteochondral plugs to ensure dimensional conformity when comparing the simulation outcomes. Agarose, a commonly used hydrogel in cartilage TE, was selected as the construct material. Its non-linear, compressible poroelastic behavior was modeled using a hyperfoam formulation with strain-dependent permeability, as in previous studies [28, 29]. Further details of the FE model development are provided in Supplementary Data S3.

The TE constructs were subjected to mechanical loading using three bioreactor setups simulating unconfined compression, combined compression-shear, and hydrostatic pressure. Boundary conditions for the first two cases were same as that for osteochondral plugs (explained in Section 2.2), except a friction coefficient of 0.1 was used between the TE construct and loading surfaces [30].

#### Dynamic hydrostatic pressure bioreactor

For knee articular cartilage *in vivo*, most of the compressive forces are transferred to hydrostatic pressure (HP) due to interstitial fluid pressurization, which acts as a primary load bearing mechanism in joints [31]. To harness this hydrostatic pressure to stimulate cartilage production in TE constructs, hydrostatic bioreactors have been used in literature imparting a wide range of loading magnitudes and frequencies [31, 32, 33]. In the present study, FE analysis of TE constructs subjected to static (0 Hz) and dynamic hydrostatic presure (at a frequency of 1 Hz) were performed. The loading magnitudes for hydrostatic pressure application were 0.5 MPa, 5 MPa, 10 MPa and 50 MPa, which were chosen to cover the wide range of magnitudes reported in literature [31]. The hydrostatic pressure was applied using a ‘Pressure Load’ in Abaqus.

## 3. Results

### 3.1. FE analysis of the human knee joint

The results of FE simulations of stance phase of gait in the human knee joint revealed that at the first peak loading (at 20% of the stance phase of gait), contact pressures were balanced between the medial and lateral compartments of the joint as observed in Figure 3. During the second peak loading (at 81.7% of the stance phase of gait), the contact pressures were higher on the lateral compartment of the tibial and femoral cartilage, suggesting a shift in load bearing towards the lateral compartment during the second peak loading of the gait cycle. This observation goes in line with previous observations in literature [17], where a gradual shift in load from the medial to the lateral compartment was observed as well. Similar trends were observed for maximum principal stress, pore pressure, pore fluid velocity and maximum shear strain as seen in Figures 3, 4. At the contact region between the femoral and tibial cartilage (Figure 8), the highest maximum principal stresses were observed in the superficial and middle zones, while maximum shear stress peaked at the cartilage surface at the point of contact. The varus-valgus rotations obtained from the simulations were within the range that was observed in experimental studies (Figure S13).

**Figure 3:**
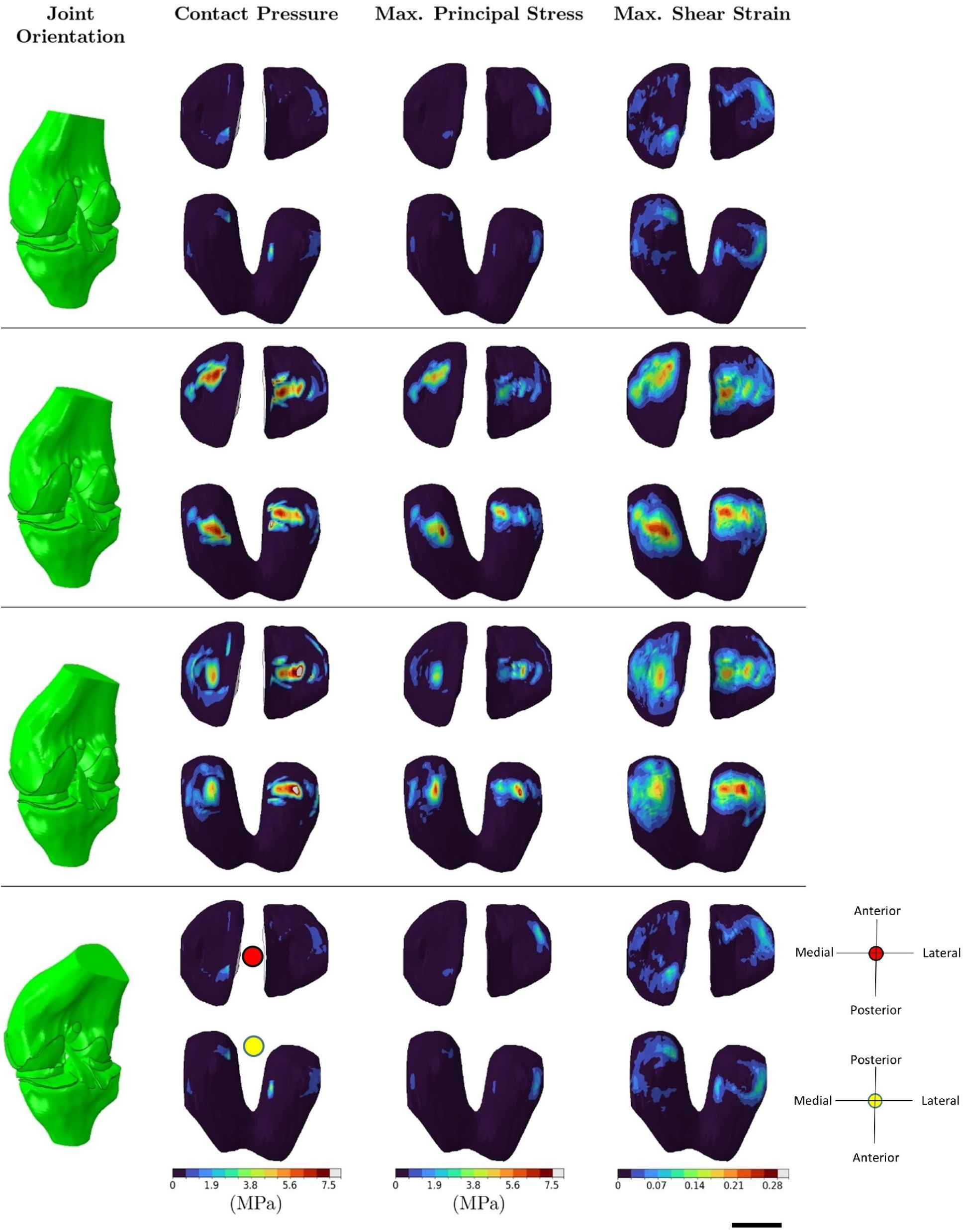
FE analysis of human knee joint during stance phase of gait showing results at fraction of stance 0, 0.2, 0.82, 1 starting from the top. Joint orientation is shown at the corresponding stance fraction. Red and yellow dots are used as indicators to show the relevant anatomical direction in context of the figures, where the tibial and femoral cartilage are displayed. Scale bar = 20 mm.

**Figure 4:**
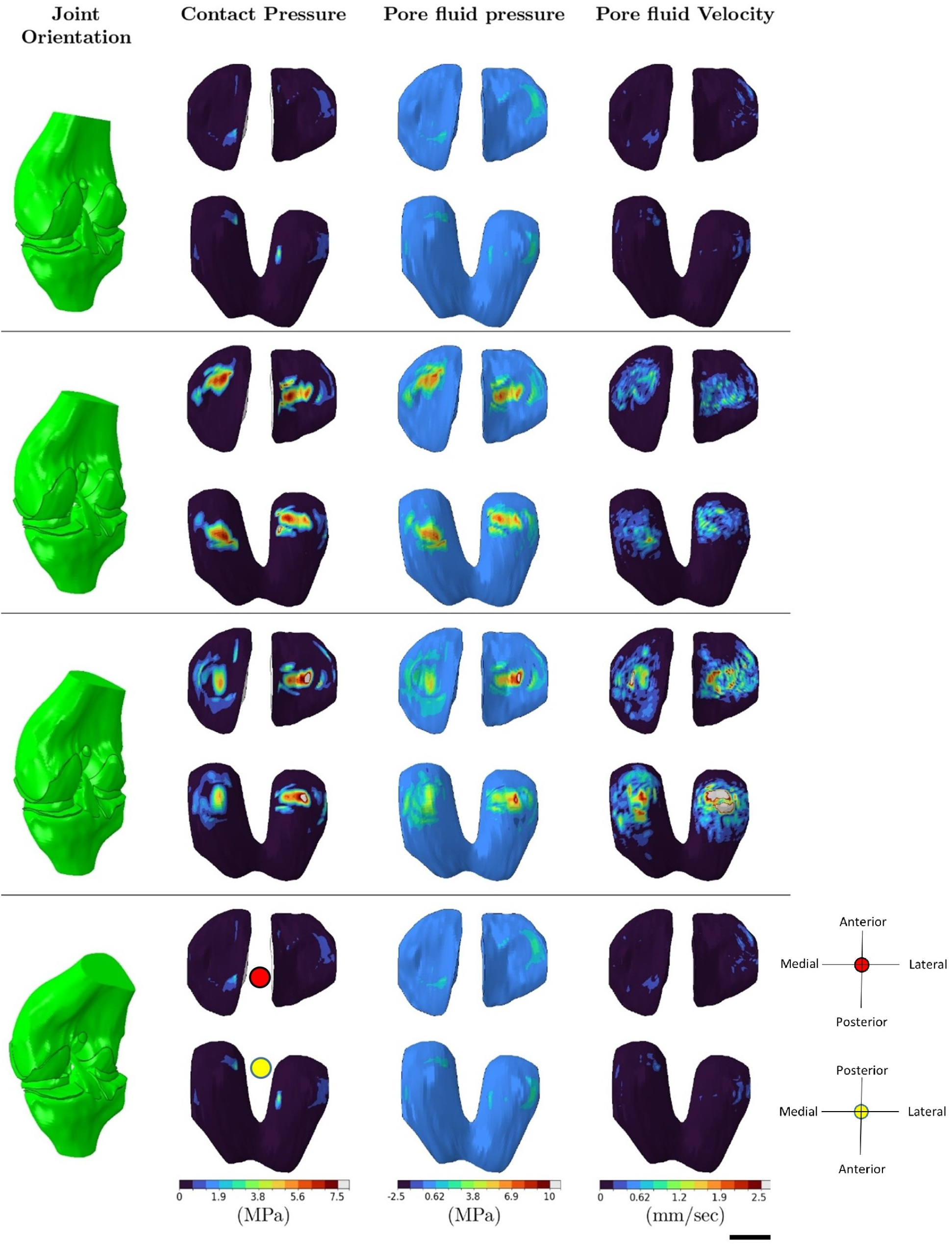
FE analysis of human knee joint during stance phase of gait showing results at fraction of stance 0, 0.2, 0.82, 1 starting from the top. Joint orientation is shown at the corresponding stance fraction. Red and yellow dots are used as indicators to show the relevant anatomical direction in context of the figures, where the tibial and femoral cartilage are displayed. Scale bar = 20 mm.

### 3.2. Role of calcified cartilage and subchondral bone in unconfined compression of osteochondral grafts

The removal of calcified cartilage from the osteochondral plug (noCC case) in simulations led to higher maximum principal stresses being transferred from the cartilage to the subchondral cortical bone for all three magnitudes of uniaxial compression (ranging from 10% to 30% strains) as compared to the intact osteochondral plug (all tissues present) (Figure 5). Additionally, there was an increase in the maximum principal stresses, pore fluid pressure, and fluid velocity around the cartilage circumference near the junction due to the absence of calcified cartilage (Figures 6, S8). For compression of only cartilage plugs, the lack of bone-cartilage interface resulted in a more uniform distribution of shear strains compared to the previous two cases (Figure S7). In the cartilage-only case, a spike in pore fluid velocity was observed at the top circumference of the cartilage, whereas, in the other two cases, the spike occurred in the cartilage circumference at the cartilage-bone interface (Figure S8). The cartilage-only case also exhibited higher overall maximum principal strains, as there was no strain transfer to the underlying tissues. Overall values of pore pressure in cartilage were consistently higher (by approx 30%) for the no CC case as compared to the full OC plugs consistently across the 3 loading regimes (Figure 6).

**Figure 5:**
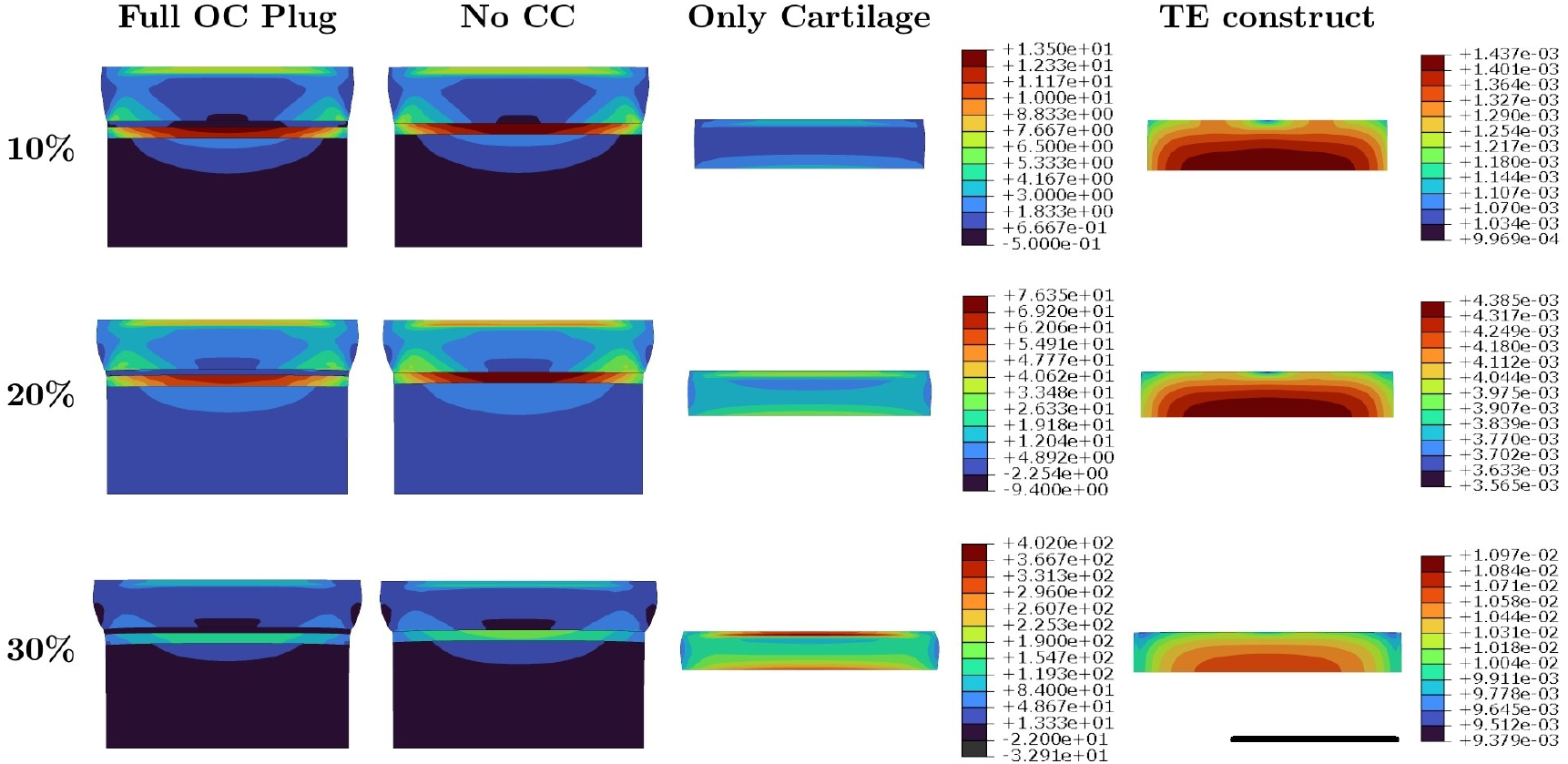
Distribution of maximum principal stress (in MPa) in different tissues of the OC plug and the TE construct for 10%, 20% and 30% strain applied in dynamic unconfined compression with 1Hz frequency. NoCC - No Calcified Cartilage. Scale bar = 5 mm.

**Figure 6:**
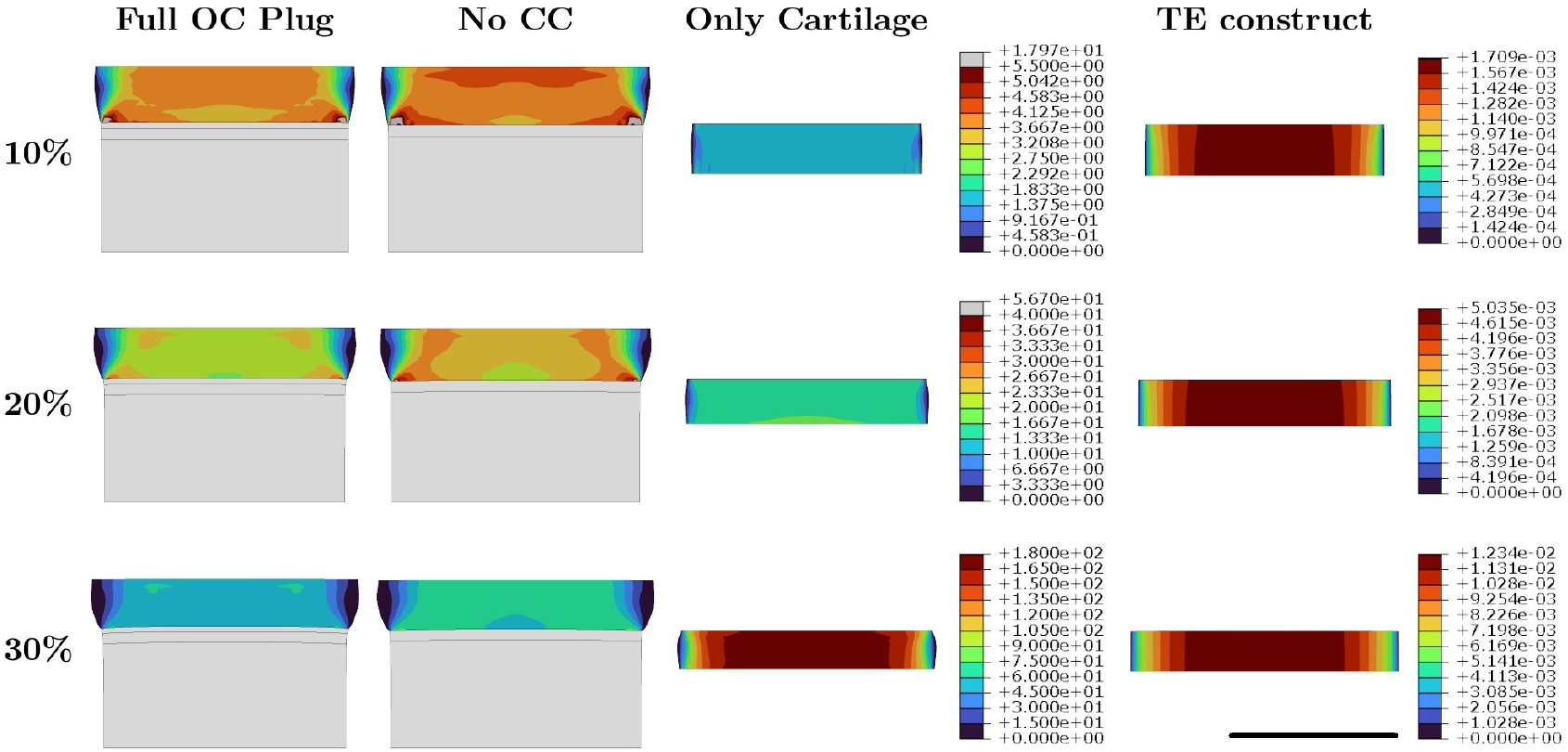
Distribution of pore fluid pressure (in MPa) in different tissues of the osteochondral (OC) plug and the tissue engineered (TE) construct for 10%, 20% and 30% strain applied in dynamic unconfined compression with 1Hz frequency. NoCC - No Calcified Cartilage. Scale bar = 5 mm.

### 3.3. Mechanical influence of combined dynamic compression and sliding rotation of ceramic ball

Dynamic compression with the ceramic ball resulted in higher maximum principal stresses at the center of the contact area in both OC plugs and TE constructs (Figure S9). In cartilage, highest stresses were observed at the SZ–MZ interface. Pore pressure was highest beneath the contact center, while fluid velocity peaked at the contact edge between the ball and the cartilage (or TE construct) (Figure S11, 7).

Rotational motion of the ceramic ball had negligible mechanical effects on cartilage. However, rotation of the ball caused asymmetric distributions of stress, pore pressure, and velocity in TE constructs, especially causing higher magnitudes in the leading direction of rotation (Figure 7, S9,S10). This asymmetry during rotation of the ball was due to the higher friction coefficient between the ceramic ball and agarose TE construct.

**Figure 7:**
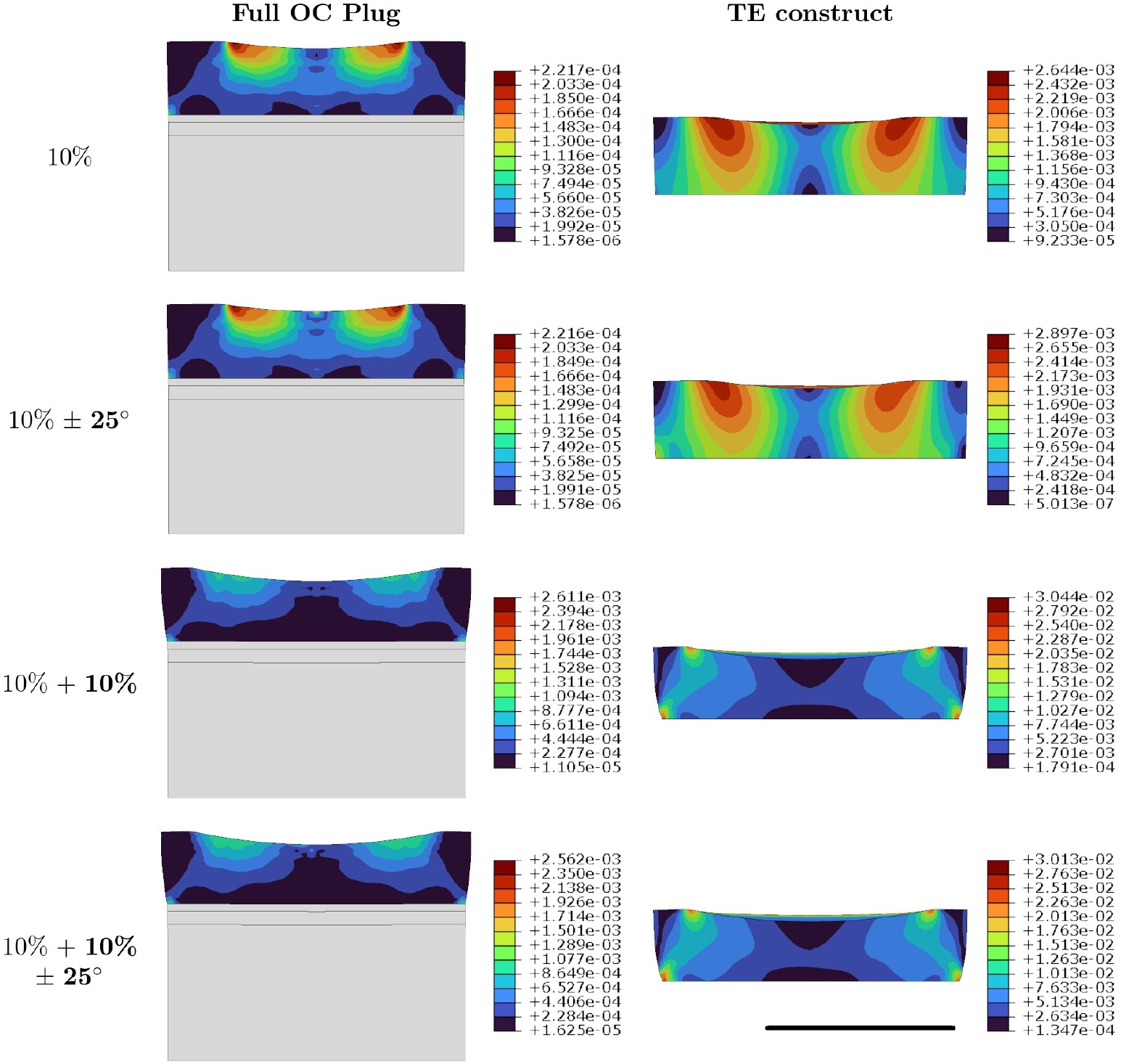
Distribution of pore fluid velocty (in mm/sec) in OC plug and the TE construct for 4 different conditions of applied compression with ball rotation. Values in bold in the first column represents a dynamic boundary condition with 1Hz frequency, while normal fonts are for quasi-static. Scale bar = 5 mm.

### 3.4. Comparative analysis of the most-physiological cases from each bioreactor type

Mechanical loading conditions from each bioreactor that most closely replicate *in vivo* knee joint cartilage mechanics during the first peak of the stance phase were selected for comparison (Figure 8). The analysis focused on four key mechanical variables: maximum principal stress, maximum shear strain, pore fluid pressure and fluid velocity. Maximum principal stress was chosen for its role in collagen fiber damage [34], maximum shear strain for proteoglycan damage [35], pore pressure for its relevance in cartilage load bearing [31], and fluid velocity for its role in loss of fixed charge density in cartilage, especially at cartilage lesions [36]. Medial tibial and femoral cartilage were used for comparison, as this compartment experiences the highest magnitudes of these variables during the first peak loading phase.

**Figure 8:**
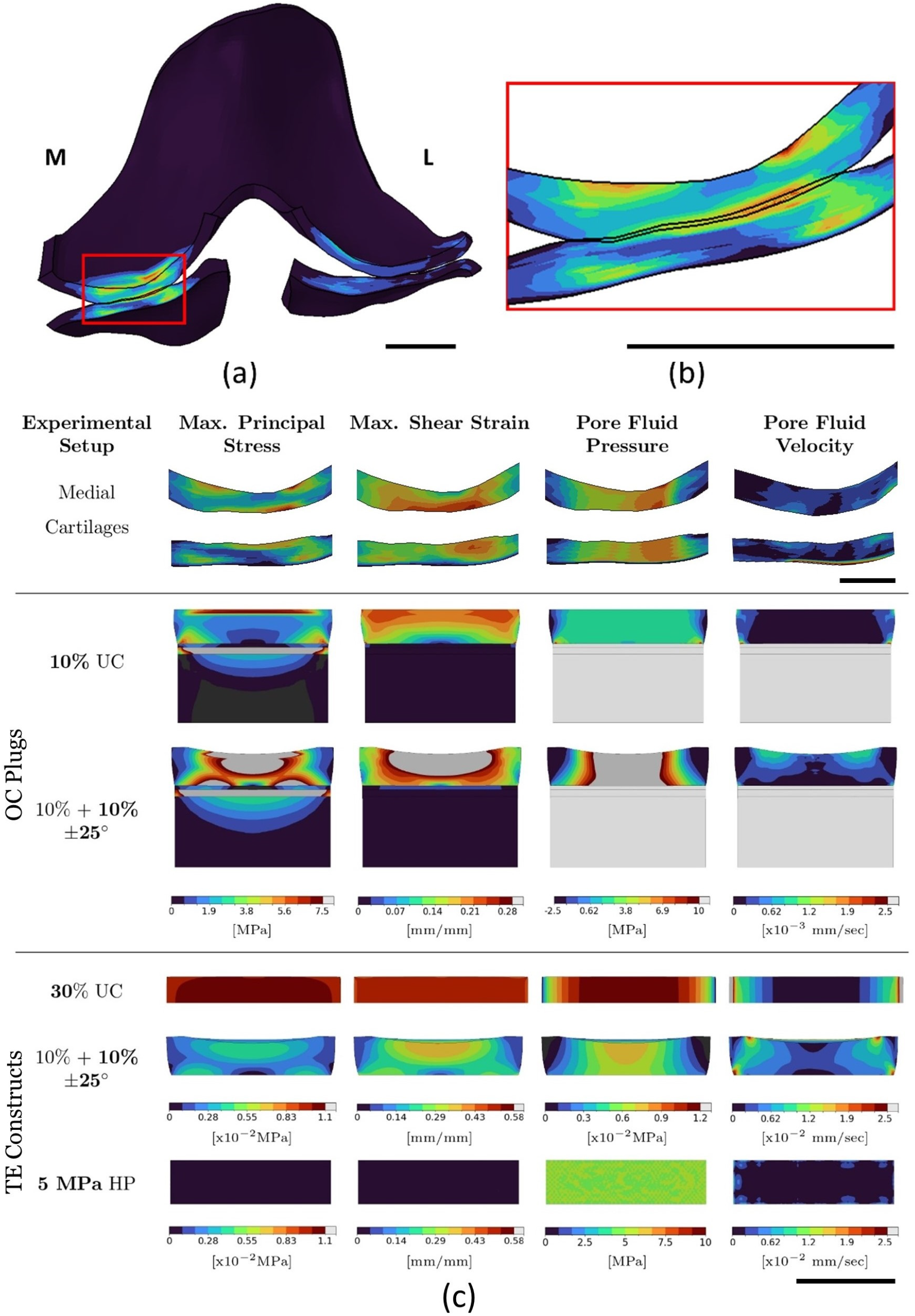
(a) Cut section of the knee joint at the first peak loading, the red box shows region of medial cartilage under contact, for which results were analyzed, Scale bar = 10 mm; (b) Zoomed in view of the contact region in (a), Scale bar = 10 mm; (c) Comparison of physical variables between the different conditions of mechanical loading of OC plugs and TE constructs with the medial cartilage region selected as in (a),(b). UC-Unconfined Compression, HP-Hydrostatic Pressure, applied at a frequency 1 Hz. Scale bar = 5 mm.

#### 10% Dynamic unconfined compression of OC plugs

In this case, the cartilage of OC plug exhibited maximum principal stress magnitudes and distributions similar to those in medial knee cartilage, with peak stresses concentrated in the superficial zone (SZ) at the interface with the middle zone (MZ). This aligns with previous studies reporting high strain concentrations and subsurface cartilage delamination at the SZ-MZ interface due to repetitive compressive loading [37, 38, 39]. Additionally, stress patterns near the cartilage–calcified cartilage interface were comparable. In both the OC plug and native cartilage, a central low-stress region beneath the contact area was flanked by zones of high stress at the bone-cartilage (or calcified cartilage) interface (Figure 8).

Maximum shear strain also followed a similar gradient in both models, decreasing from the surface to the deeper zones. Unlike principal stress, maximum shear strain was not concentrated in the SZ but was more evenly distributed. While principal stress is influenced by tension in surface-aligned collagen fibers, shear strain depends on the shear modulus of proteoglycan-rich matrix components. This difference in mechanisms of load bearing of the cartilage constituents might explain the difference in distribution of the stress components.

Pore pressure distributions were also similar, approaching zero near the free-draining surfaces in both the knee and OC plug. However, pore pressure magnitude was lower in the OC plug. Pore fluid velocity was highest near the contact area in knee cartilage. In the OC plug, peak fluid velocities occurred along the curved free-draining surface, where a zero pore pressure boundary condition was applied.

#### 30% Dynamic unconfined compression of TE constructs

Due to the low stiffness of agarose, stresses in the TE construct were 2 orders of magnitude lower than in knee cartilage, even under 30% compression. Maximum principal stress and shear strain were uniformly distributed across the construct thickness, unlike knee cartilage where SZ stresses are higher due to collagen fiber alignment. Maximum shear strain was twice that of knee medial cartilages. Fluid velocity was 10 times higher and pore pressure 100 times lower than in knee cartilage, though their spatial distributions were similar.

#### Dynamic compression and sliding motion applied to OC plugs

A quasistatic compression of 10% followed by a 10% dynamic compression and simultaneous *±*25^°^ dynamic rotation of the ball was chosen for comparison with the knee articular cartilage. The values of all the 4 mechanical variables obtained from this bioreactor condition were higher in magnitude than the ones observed for knee cartilage (Figure 8). High maximum principal stress and pore fluid pressure distributions were observed at a region in the center of cartilage under the contact area of the ball with the cartilage, similar to the area underneath cartilage to cartilage contact in the knee joint. Highest values of maximum principal stress and maximum shear strain were located in the SZ. Pore fluid velocity in the OC plug followed similar distribution and magnitude, with highest magnitudes occurring towards the edge of the contact (between the ball and cartilage) and close to the top surface, similar to the knee cartilage.

#### Dynamic compression and sliding motion applied to TE constructs

For the same 10% dynamic compression and *±*25^°^ sliding motion as in the previous case, maximum shear strain close to the TE construct surface was significantly higher (around double) than that observed for knee joint cartilage. Distribution of both maximum principal stress and maximum shear strain followed a gradient across the construct thickness with higher values closer to the contacting surface, similar to what is observed in knee articular cartilage. The magnitudes of pore fluid pressure and velocity followed similar trends as observed for the unconfined compression of TE constructs as described above.

#### Dynamic Hydrostatic Pressure on TE Constructs

Applying 5 MPa dynamic hydrostatic pressure produced a uniform pore pressure distribution matching that of knee cartilage beneath contact areas. This was the only setup that successfully mimicked physiological pore pressure in TE constructs despite their high permeability. However, other mechanical variables remained significantly lower than in native cartilage, highlighting the limitation of this method in replicating full mechanical behavior.

## 4. Discussion

The present study compared common bioreactor setups used to apply mechanical stimuli to osteochondral (OC) explants and cartilage tissue-engineered (TE) constructs against a finite element (FE) model of the human knee joint simulating stance phase of gait. While prior studies have examined mechanics of individual bioreactors [40, 11, 8, 10], this work uniquely benchmarks them against an *in vivo* physiological reference, enabling a more informed assessment of how closely each setup replicates native joint mechanics.

The study revealed that no single bioreactor replicated all mechanical variables in both magnitude and spatial distribution. However, the findings offer guidance for selecting bioreactors based on specific research goals. For instance, 10% unconfined compression of OC plugs matched native cartilage in terms of stress and pressure, but fluid velocity patterns differed. In TE constructs, unconfined compression (up to 30%) produced stresses and pressures 2–3 orders of magnitude lower than in knee cartilage. Hydrostatic loading at 5 MPa better approximated native pore pressures, though it induced much lower strains.

The 8 mm diameter of the OC plugs was chosen to match the typical cartilage contact area (6–10 mm) during peak gait loading. Unlike *in vivo* cartilage, which is laterally constrained by surrounding tissue, the OC plugs had free lateral surfaces that allowed unrestricted bulging and fluid outflow. Consequently, unconfined compression led to higher fluid velocities at the plug’s lateral surface. This ‘edge effect’ could have led to the loss of matrix constituents from the lateral surface due to fluid flow as observed in some *in vitro* studies [23], especially since harvesting of the plugs by drilling it from the joints would have already caused damage to edges of the OC plug. For the combined compression and ball rotation simulations, though the effect of ball rotation on cartilage mechanics was negligible, it was still considered for the comparative study since ball rotation mimics the sliding motion between the knee articular cartilage during gait, which is known to trigger lubricin production in the SZ of cartilage. Lubricin is responsible for maintaining the low friction coefficients of cartilage [26].

This study also elucidated the importance of calcified cartilage layer in firstly, reducing stress transfer to the underlying subchondral bone during compression, and secondly, acting as an intermediate layer between bone and cartilage to reduce interfacial stresses in cartilage. Calcified cartilage in the study was modeled as a linear elastic material, since permeability of calcified cartilage is not well-defined experimentally. Some studies in the literature have characterized it as entirely impermeable, while others have reported its permeability to be lower or comparable to that of articular cartilage [41]. In future studies, experimentally determined permeability of calcified cartilage should be incorporated into the poroelastic FE analysis of OC grafts to simulate fluid (and thereby molecular) cross-talk between the cartilage and bone through the calcified cartilage. This cross-talk is expected to influence and potentially alter the progression of cartilage degeneration in osteoarthritis [42].

The FE analysis of mechanical stimulation of agarose TE constructs in different bioreactors highlighted that lower maximum principal stresses were developed in the constructs due to the relatively compliant behavior of hydrogel as compared to articular cartilage. Furthermore, the TE constructs were homogeneous without any anisotropy, which resulted in a more uniform distribution of stresses as compared to cartilage where the anisotropy of collagen fibrils led to higher stresses in the SZ. The FE analysis also revealed that TE constructs developed lower pore pressures and higher pore fluid velocities for the same magnitudes of mechanical loading as compared to cartilage. The lower pore pressure meant lower load bearing capacity of the construct, which would restrict its application when implanted *in vivo* in a cartilage defect. High pore fluid velocities can have a two-fold effect. On the one hand, it might facilitate better distribution of the nutrient rich medium throughout the construct, thereby favoring cartilage regeneration [43]. On the other hand, high fluid flows might disrupt the nascent matrix secreted by the embedded cells in the hydrogel, thereby leading to matrix loss and weakening of the construct as elucidated in a study by Orozco et al. [44].

In the OC plugs, the cartilage was attached rigidly to the underlying calcified cartilage and subchondral bone. This constraint resulted in stress concentration at the interfacial region for cartilage. However, for the unconfined compression of TE constructs, free lateral expansion of the construct was possible, thereby altering the deformation field as compared to native cartilage. To prevent this from happening, and also to reduce fluid flow from the lateral sides of the construct, partial confinement of half the thickness of the construct from the bottom had been suggested in a previous study in literature by Thorpe et al. [45]. Furthermore, restrictions imposed by the partial confinement on fluid flow and lateral deformations would result in higher pore pressures and also would create a gradient of the mechanical environment that would more closely mimic the *in vivo* mechanics of cartilage.

Finally, in the current study, mechanics of articular cartilage and TE constructs were studied at the macro-scale, tissue-level resolution. This approach provided valuable insights into the distribution of physical variables across different setups and under varying loading conditions. However, the present study did not consider any cell-matrix interactions, which is known to influence behavior of cells to mechanical stimuli [46]. In the future, efforts must be made to develop multiscale models, that links the tissue-level mechanics (as done in this study) with micro-scale cellular mechanics, and subsequently the intracellular mechanotransduction processes that are triggered as a result of mechanical stimuli sensed by individual cells which are located at different regions of the cartilage or TE construct. Such a multiscale modeling framework would provide more granularity while quantifying loading protocols across different bioreactor setups to facilitate tissue regeneration. To conclude, this study provides a robust framework for comparing bioreactor loading conditions with *in vivo* joint mechanics, offering valuable insights for optimizing cartilage tissue engineering strategies. By bridging the gap between experimental and physiological conditions, the findings will contribute to improving bioreactor design, enhancing regenerative medicine approaches, and advancing treatments for cartilage repair.

## 5. Acknowledgments

This study received funding from the European Union’s Horizon 2020 research and innovation programme under the Marie Skłodowska-Curie grant agreement No. 721432 (CarBon project), the In Silico World project (grant agreement No. 101016503), the European Research Council Consolidator Grant No. 101088919 and the Belgian Federal Public Service Policy & Support (grant DigiTwin4PH). The funding sources have no role in design and execution of the study.

## 6. Conflict of interest statement

Wouter Wilson is an employee of Sioux Technologies, Eindhoven, Netherlands.

## Appendix A. My Appendix

Supplementary data and figures are found in the corresponding Supplementary Material.

## Supplementary Material

### S1. FE modeling of the human knee joint

#### S1.1. Geometry and Meshing

The geometry of the tibiofemoral joint of a cadaveric right knee from a 70 year old female donor (77.1 kg, 1.68 m) was obtained from the Open Knee database [1] and imported in Abaqus 2023 (SIMULIA, Providence, RI, USA). The Open Knee joint geometry included geometric representations of the bones (femur and tibia), cartilage(femoral and tibial), menisci (medial and lateral), and ligaments (anterior and posterior cruciate, medial and lateral collateral). The segmented geometry of the lateral collateral ligaments in the database was incomplete. So, the collateral ligaments (both medial and lateral) were excluded from the simulations. Since the collateral ligaments are primarily responsible for side-to-side stability of the knee joint, and do not contribute to load-bearing in the joint, their exclusion are expected to not significantly alter the stresses in articular cartilage. The anterior and posterior ends of the menisci were attached to the tibial plateau by means of a series of linear springs with total stiffness of each meniscal attachment (also called meniscal horns) set to a 2000 N/mm based on previous studies [2, 3]. The femoral and tibial cartilage were rigidly attached to the femur and tibia respectively using the ‘Tie’ constraint.

Articular cartilage were meshed with hexahedral pore pressure elements(C3D8P) and the other soft tissues were meshed with hexahedral linear elements (C3D8). The tibia and femur were meshed with rigid elements (R3D4). A mesh convergence study was conducted by varying the number of elements across the thickness of the articular cartilage from 7 to 12. These numbers were based on previous studies in literature that used similar fibril-reinforced poroviscoelastic (FRPVE) material model for knee joint, where the number of elements across the cartilage thickness ranged from 4-6 [3]. This adjustment resulted in a total element count ranging from 60,837 to 104,292 for the coarser and finer meshes, respectively. No significant changes were observed in the distribution of the maximum principal stresses, pore pressures and contact pressures from the coarser to finer mesh as can be seen from Supplementary Figure S4.

**Figure S1:**
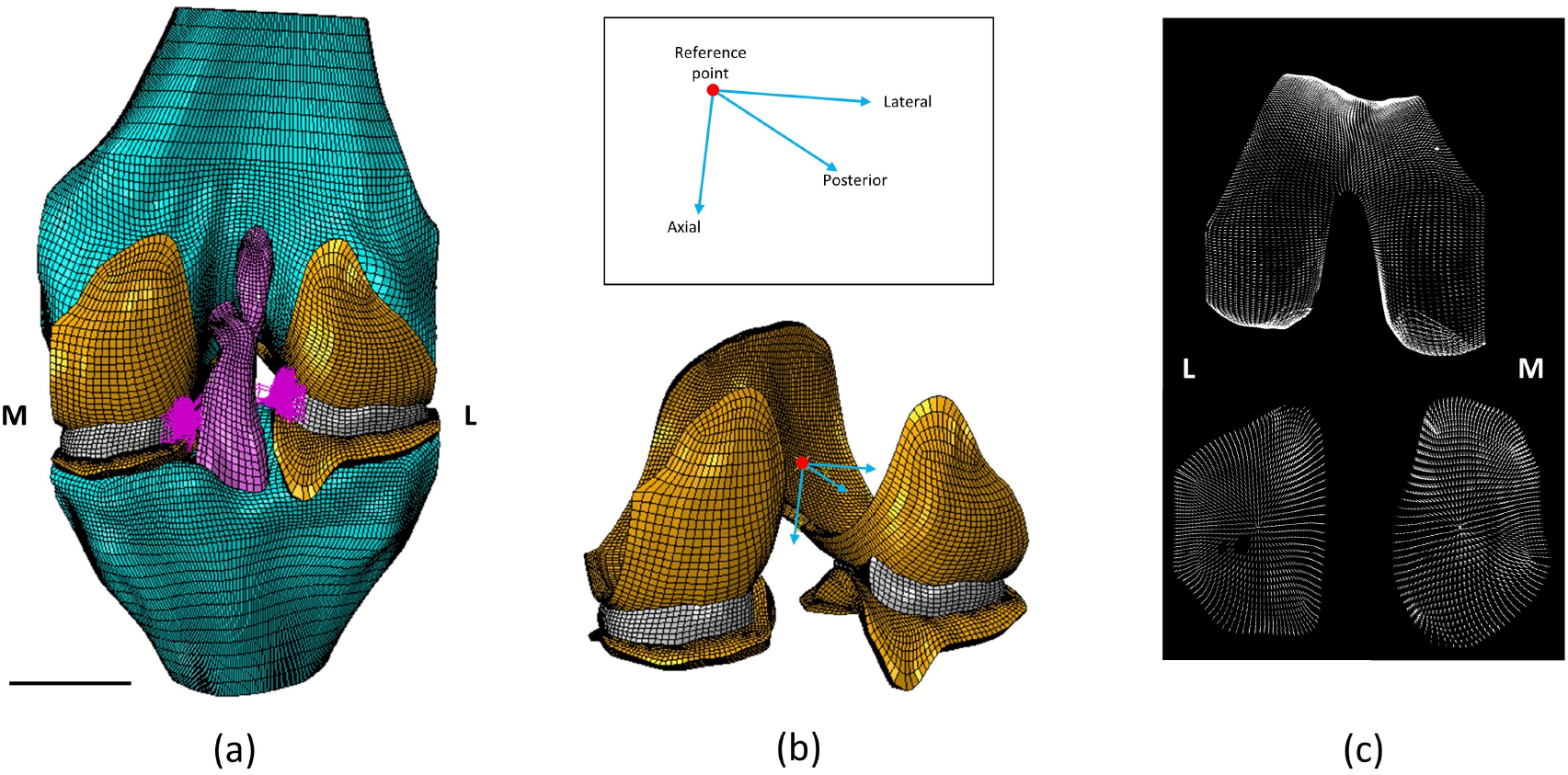
(a) FE model of the human knee joint with the different tissues and meniscal attachment springs (in pink), M-medial, L-lateral, Scale bar = 20 mm; (b) The reference point and the axes about which boundary conditions are applied for the FE model (ligaments and bones have been removed for visualization). In the inset box, a zoomed in view showing the reference point and axes with anatomical directions; (c) Orientation of the split-line patterns at the superficial zone as implemented in the FE model.

**Figure S2:**
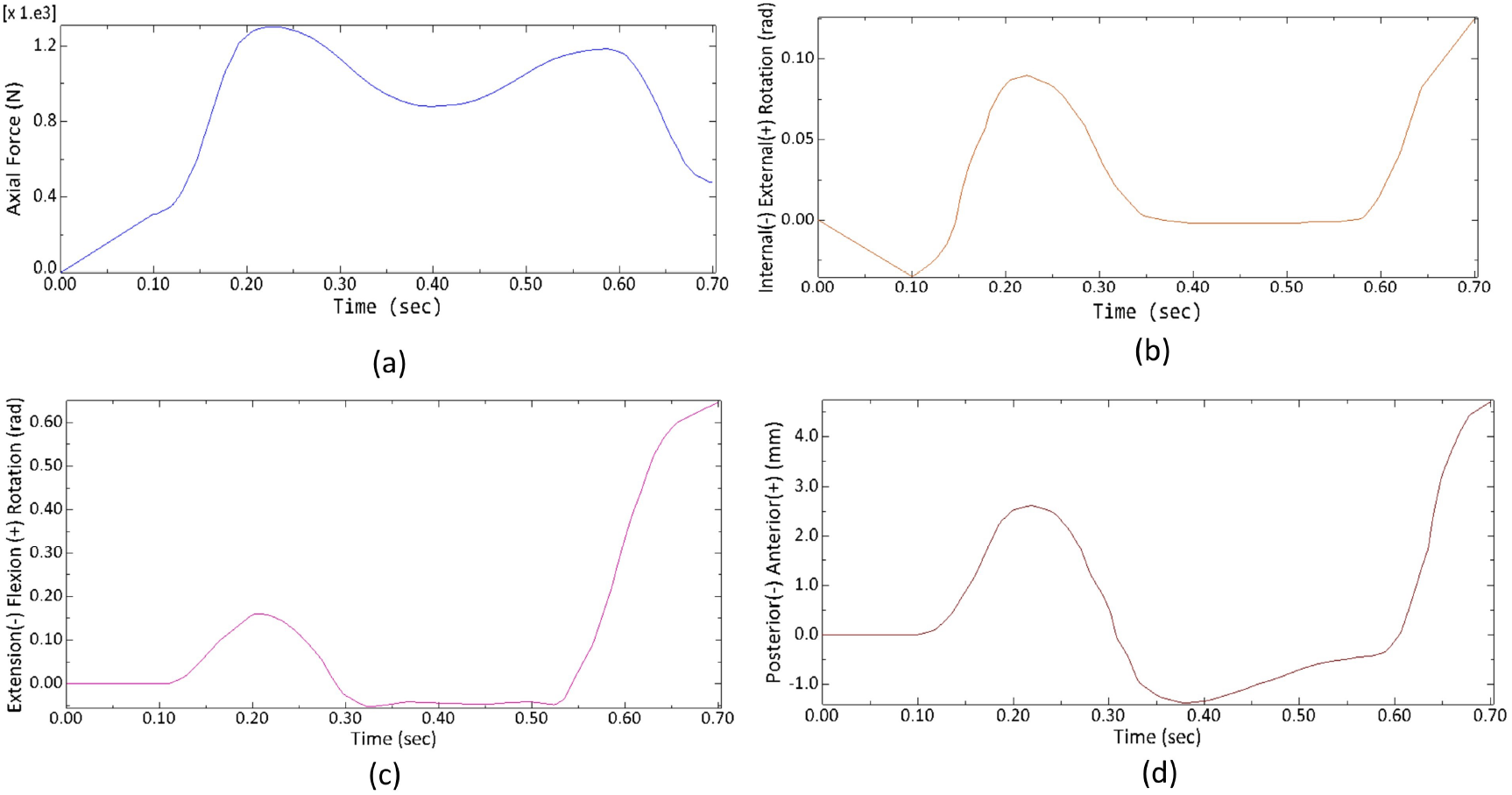
Boundary conditions applied to FE model of the human knee joint as adjusted from [3]. (a) Axial compressive force (N). (b) Internal-external rotation (radians). (c) Extension-flexion rotation (radians). (d) Posterior-anterior translation (mm).

#### S1.2. Constitutive model of tissues

Articular cartilage was modeled as a fibril-reinforced poroviscoelastic material and implemented using UMAT subtroutine, as was described in previous studies [4, 5]. This biphasic material model consists of a solid and pore-fluid component, where the solid component consists of a fibrillar and non-fibrillar part. The fibrillar part models the collagen fibrils present in articular cartilage, whereas the non-fibrillar part mainly comprises proteoglycans. The total stress at an integration point was given by:

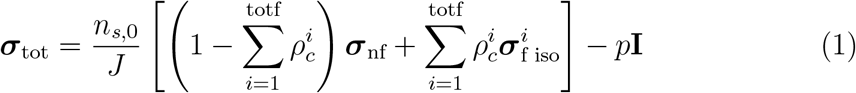

where **I** is the unit tensor, *n*_*s*,0_ is initial solid volume fraction, *J* is the determinant of the deformation tensor **F**, totf is the total number of fibrils considered at each integration point, 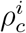 is the volume fraction of the *i*^*th*^ collagen fibrils with respect to the total volume of the solid matrix, ***σ***_nf_ is the stress in the non-fibrillar matrix, ***σ***_f iso_ is the stress in the collagen fiber network.

The strain-dependent permeability of cartilage was given by

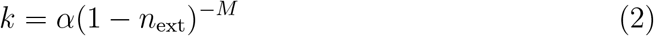

where *α* and *M* are positive material constants.

##### Non-fibrillar Part

The non-fibrillar part of cartilage, which primarily consists of proteoglycans was modeled using a modified Neo-Hookean law as in Heuijerjans et al.[6]. The stress in the non-fibrillar matrix ***σ***_nf_ was calculated as:

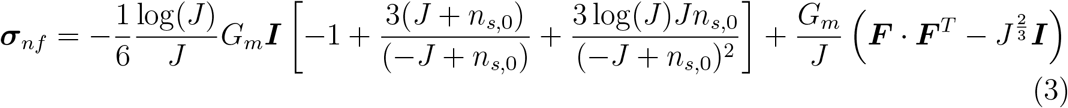

where *G*_*m*_ is the shear modulus of the non-fibrillar part.

##### Fibrillar Part

The fibillar component consisted of 2 primary and 24 secondary collagen fibrils at each integration point. The primary fibrils followed an arcade-like structure along the cartilage depth, with the fibrils perpendicular to the bone-cartilage interface at the deep zone and parallel to the cartilage surface in the superficial zone of cartilage. The split-line patterns of the collagen fibrils in the femoral and tibial cartilage were implemented by using in-house codes developed in Matlab R2022b (The MathWorks Inc., Natick, Massachusetts)(Figure S1(c)). The stress in each fibril consisted of an isotropic component ***σ***_iso_ that accounts for the isotropic stiffness of the collagen fibers and a stretch activated component ***σ***_f_ along the fibril direction. The isotropic component of stress was calculated using the same Neo-Hookean material model as used for the non-fibrillar part, with the shear modulus of collagen fibers incorporated *Gm*_*f*_ . The stretch-activated directional component of the stress was calculated assuming a spring (S1) in parallel to a spring (S2) and dashpot (*η*) in series. The equations for the fibril stresses are as described below:

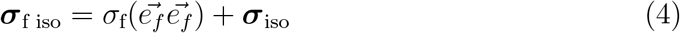

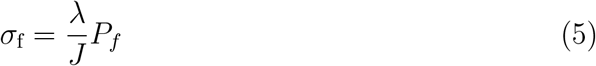

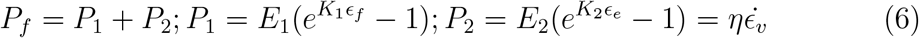

where, *P*_*f*_ is the total fibril stress, 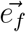 is unit vector along the current fibril direction, *ϵ*_*f*_ is the total fibril strain, *ϵ*_*e*_ is the strain in spring S2 and *ϵ*_*v*_ is the strain in the dashpot. *E*_1_, *E*_2_, *K*_1_ and *K*_2_ are material constants, *λ* is elongation of the fibril. The density of each collagen fibrils is given by

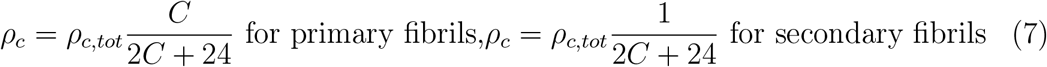

where, *C* is a positive constant and *ρ*_*c,tot*_ is the total collagen fiber density.

The fluid fraction (*n*_*f*_), collagen fraction (*n*_*coll*_) and fixed charge density (*c*_*F*_) were distributed as a function of the normalized depth *z*^*^ [7] as shown below:

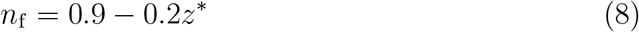

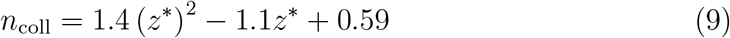

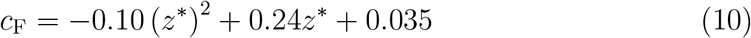

Table S1 lists the values of the material parameters that were used for the simulations.

**Table S1:**
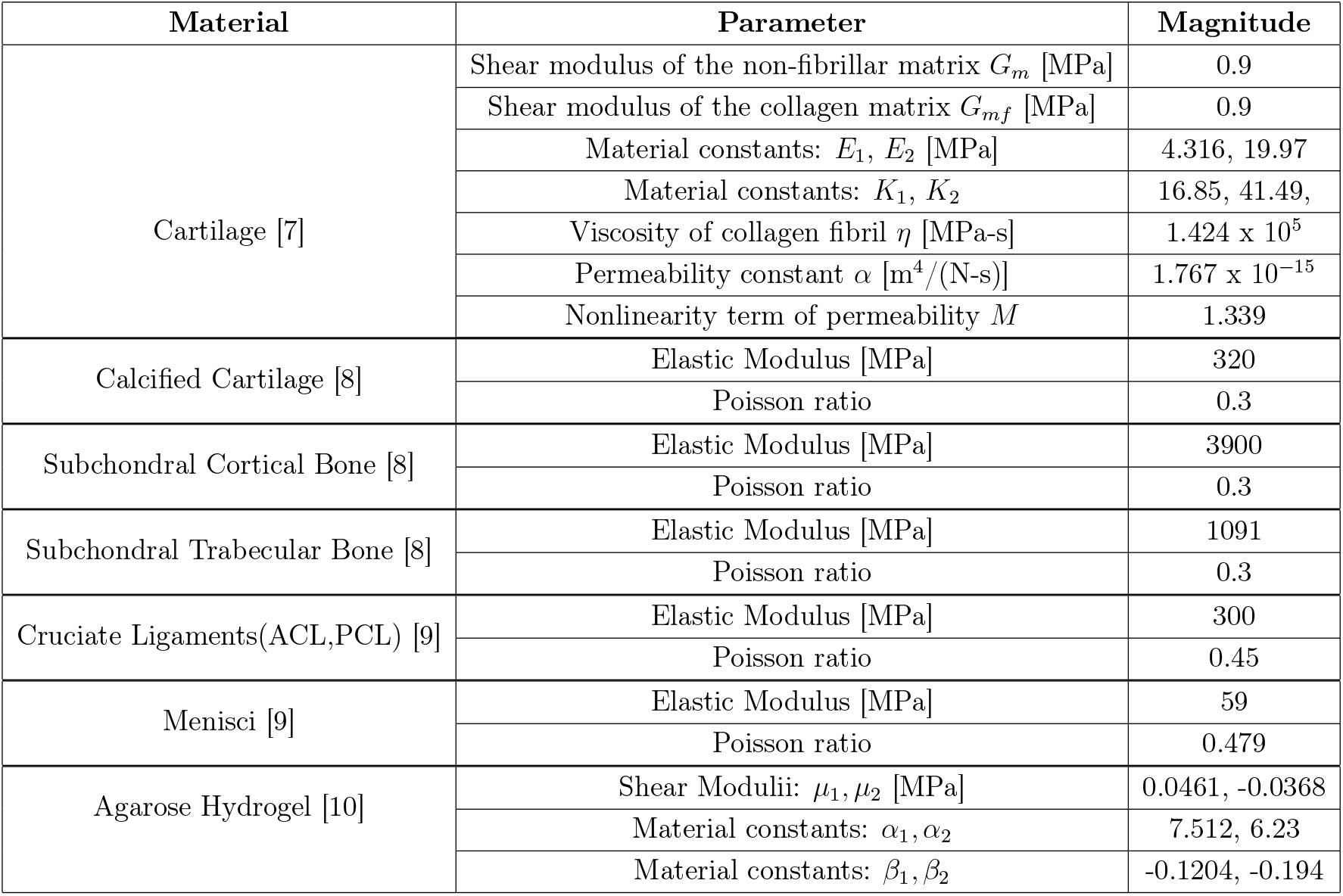
Table to describe the material parameters of the tissues and hydrogel used in the study.

#### S1.3. Boundary Conditions and Contact Interactions

Time-dependent translations, rotations and forces were implemented as boundary conditions to the femur to simulate the stance phase of the gait cycle as in Figure S2. The magnitude of joint axial force was scaled by a factor of 0.77 from the original values obtained from [3] to account for patient specific body weight. The boundary conditions were applied to a reference point located at the center of the medial and lateral femoral epicondyles as shown in Figure S1(b). The tibia was fixed in all directions. The gait cycle was simulated in the FE model in two successive steps as discussed below:

##### Initial step

Since the segmented geometry of the knee was at full extension, an initial step was performed to adjust the initial axial force and rotations (internal–external and extension–flexion) to match the values at 0% of the gait cycle. Translations were maintained at their initial positions. The initial step was assigned a duration of 0.1 seconds.

##### Gait cycle step

In this step, femoral translations (anterior-posterior) and rotations (internal–external and extension–flexion) were applied as time-dependent boundary conditions at the reference point using the smooth step option in Abaqus. The varus-valgus rotation was kept free to ensure continuity of contact between the femoral and tibial cartilage throughout the entire stance phase of the gait cycle. Also, by comparing the resulting varus-valgus rotation from simulations with respect to experimentally observed values, the role of supporting tissues (ligaments) in governing the passive motion of the knee joint during the stance phase could be elucidated. The medial-lateral translation was fixed to prevent the menisci from sliding out and losing contact with tibial cartilage in absence of collateral ligaments, which could lead to errors in contact convergence. Axial force variation during the stance phase was applied as a concentrated force at the reference point. The duration of a single stance was set to 0.6 seconds [11].

##### Contact definition

The general contact algorithm with selected surface pairs was used in Abaqus to define contact between the femoral and tibial cartilage, and the menisci. Frictionless behavior was assumed for the contacting surfaces. For the normal behavior, the penalty contact constraint enforcement method was used. The penalty stiffness was scaled by a factor of five times the default value, to minimize the interpenetration between contacting surfaces of the femoral and tibial cartilage.

### S2. FE modeling of bioreactors for osteochondral plugs

#### S2.1. Geometry and Meshing

Three dimensional (3D) geometries of cylindrical osteochondral plugs were created in Abaqus. A diameter of 8 mm was chosen for the plugs based on studies in literature [12, 13]. The osteochondral plugs consisted of cartilage (2 mm thick considering average human cartilage thickness), calcified cartilage (0.2 mm thick [14]) and 4 mm of subchondral bone underneath. The subchondral bone was further subdivided into subchondral cortical bone (0.36 mm thick) and subchondral trabecular bone (3.64 mm thick) [15]). To assess the role of individual tissues in the stress response of osteochondral plugs, three cases were simulated by progressively excluding specific tissues (Figure S3):

a. All: In this case all the constituent tissues as described above were considered.
b. No calcified cartilage (NoCC): The calcified cartilage was removed from the plugs, with all other tissues intact, to study the effect of calcified cartilage in load bearing.
c. Only cartilage: This case consisted of only the cartilage of 2 mm thickness.

Cartilage was meshed with hexahedral pore pressure elements (C3D8P) while calcified cartilage and subchondral bone were meshed with hexahedral elements (C3D8). A mesh convergence study was performed to determine the final mesh density for the simulations (Supplementary Figure S5)

**Figure S3:**
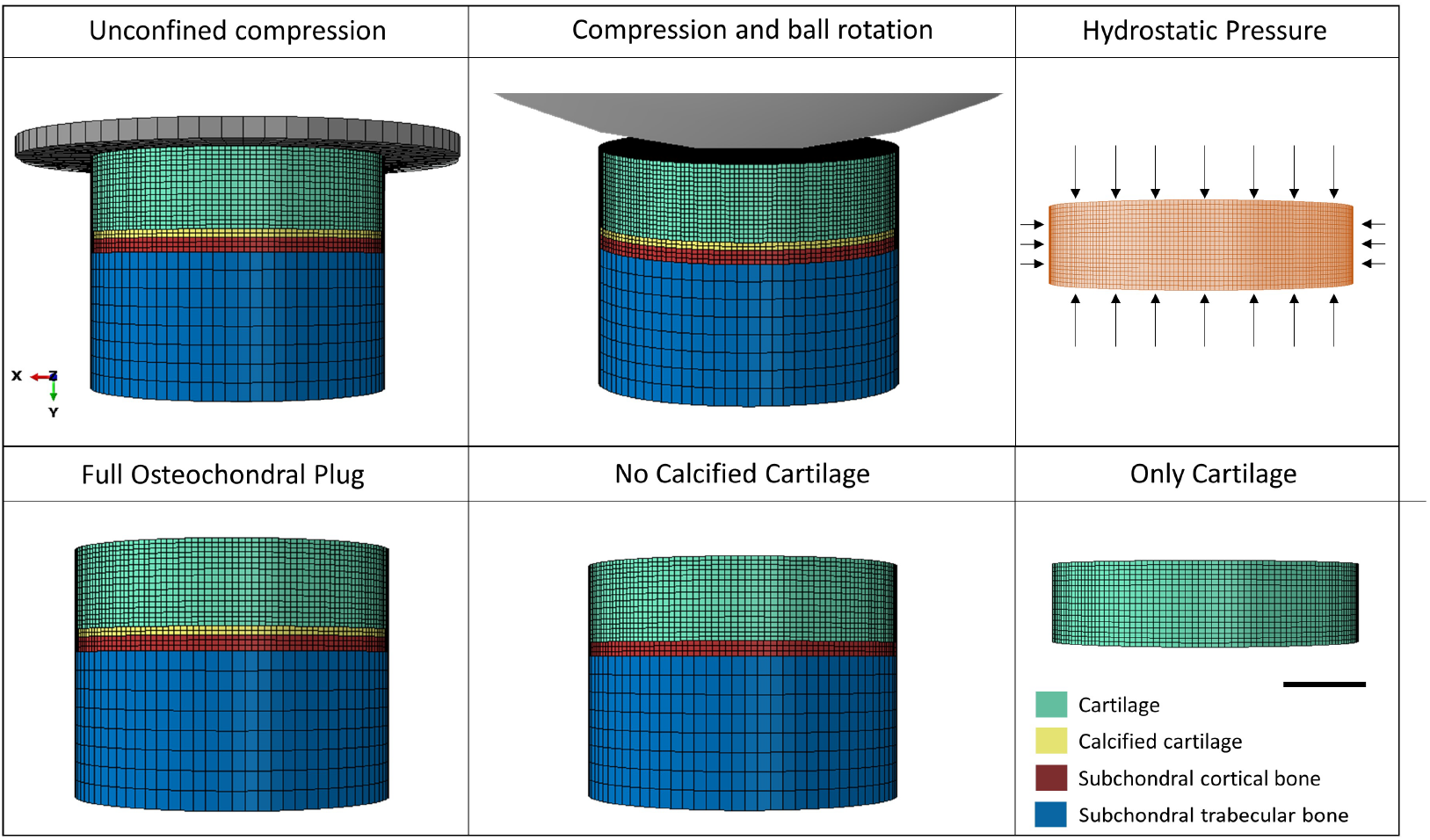
Top row: Different types of bioreactors used in the study. The first two from left show osteochondral plugs, while hydrostatic pressure was applied to only TE constructs. A portion of the ball is visualized in the compression and ball rotation case. Bottom row: FE models to study the role of different tissues in osteochondral plugs. Scale bar = 2 mm.

#### S2.2. Constitutive model of tissues

Cartilage was modeled as a fibril-reinforced poroviscoelastic material as described in the previous section. Calcified cartilage, subchondral cortical and tra-becular bone were modeled as linear elastic materials with material parameters obtained from literature [8], and mentioned in Table S1.

#### S2.3. Boundary Conditions and Contact Interactions

The osteochondral plugs were subject to mechanical loading in three different bioreactor setups that are commonly used in literature to apply unconfined compression, confined compression and combined compression and shear respectively. The boundary conditions that were applied to mimic mechanical loading in FE model of the bioreactors are described below (Figure S3) :

##### (a) Dynamic unconfined uniaxial compression bioreactor

This bioreactor is most commonly used in literature for its simplicity in design and ease of application of mechanical loading [16, 17]. Uniaxial dynamic compression was applied to the plugs in the FE model by using a rigid plate with a diameter of 12 mm. A haversine function with a frequency of 1 Hz and amplitudes of 10%, 20%, and 30% of the initial cartilage thickness was defined at the reference point of the rigid plate to impart dynamic compression to the plugs. The 1 Hz frequency closely resembles that of human gait. The loading amplitudes were selected to encompass the range of hypo to hyper-physiological loading regimes mentioned in literature [16, 18]. Vertical displacement of the the bottom surface of the plugs was constrained using the ‘Ysymm’ boundary condition in Abaqus. Furthermore, to eliminate the possibility of any rigid rotations or translations of the plugs in lateral direction along the X-Z plane, two diametrically opposite nodes at the bottom surface of the plug were fixed in both the X and Z directions, respectively. This approach prevented rigid motions caused by numerical approximations without affecting the mechanics of uniaxial unconfined compression. To allow free fluid flow across the curved surface of the cartilage, a zero pore pressure boundary condition was prescribed at the nodes on that surface. A surface-to-surface contact was defined between the rigid plate and the top surface of cartilage with a ‘hard’ normal pressure-overclosure behavior. A friction coefficient of 0.003 [19] was applied between the contacting surfaces using the ‘Lagrange Multiplier’ friction formulation. For unaxial compression, all the 3 types of plugs were considered (all, no calcified cartilage and only cartilage) to do a comparative study of the contribution of different tissues to mechanical stress response. The corresponding tissues were rigidly attached to one another by using a ‘Tie constraint’.

##### (b) Dynamic multi-axial loading bioreactor

Articular cartilage in the human knee joint is subject to a multi-axial state of loading during joint movements. To replicate this multi-axial loading in experimental settings, bioreactors were designed to apply a combination of shear and compressive forces to osteochondral plugs. One such bioreactor, as developed in studies from AO Davos, Switzerland [20, 21] used a commercially available ceramic hip ball (32 mm in diameter) to impart multi-axial loading. This was achieved by simultaneously applying axial compression through displacement and shear force by rotation of the ball about an axis perpendicular to the direction of axial compression. In the FE model of this setup, the ceramic ball was modeled as a rigid body due to its high stiffness as compared to cartilage. Four loading conditions were tested in this setup, with the boundary conditions applied to a reference point at the center of the ball: (1) 10% quasi-static compression for 10 seconds, (2) 10% quasi-static compression, followed by 10% dynamic loading at 1 Hz, (3) 10% quasi-static compression, followed by ±25 degree dynamic rotation at 1 Hz, (4) 10% quasi-static compression, followed by 10% dynamic loading and ±25 degree dynamic rotation at 1 Hz. For the FE analysis of the three different types of bioreactors discussed above, dynamic loading was applied for five cycles. The results were then analyzed at the peak loading of the fifth cycle to ensure that the initial transient effects had dissipated.

### S3. FE modeling of bioreactors for cartilage tissue engineered constructs

#### S3.1. Geometry and Meshing

The tissue-engineered (TE) constructs were modeled as discs of 8 mm diameter and 2 mm thickness. The diameter and thickness were kept consistent with those of the cartilage in osteochondral plugs to ensure dimensional conformity when comparing the simulation outcomes. The TE discs were meshed with hexahedral pore pressure elements (C3D8P). A mesh convergence study was performed to obtain the final mesh density as shown in Supplementary Figure S6.

#### S3.2. Constitutive model of TE construct

The TE constructs were assumed to be made of agarose, a hydrogel which is frequently used for cartilage TE in literature. Since agarose is known to behave as a compressible non-linear poroelastic material [22], a hyperfoam model with strain-dependent permeability was used to describe its nonlinear poroelastic behaviour as done in previous studies [10]. For the hyperfoam behavior, a modified Ogden-Hill strain energy potential (hyperfoam) available in Abaqus was used as given by the constitutive relation below:

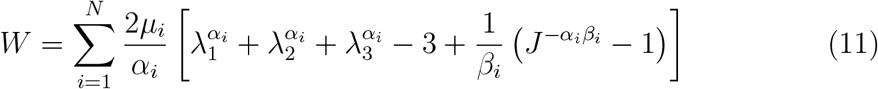

where *W* is the strain energy density function and *µ*_*i*_, *α*_*i*_, and *β*_*i*_ are material constants. *J* is the determinant of the deformation gradient tensor and the principal stretches are given by *λ*_1_, *λ*_2_, *λ*_3_. *N* for agarose is *N* = 2.

Strain-dependent permeability for agarose was implemented as in Khoshgoftar et al. [10]:

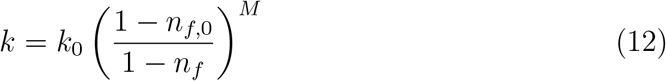

where *k*_0_ is the initial permeability, *M* is a positive constant, and *n*_*f*,0_ and *n*_*f*_ are the initial and current fluid fractions respectively. Material parameters used for the simulations for 2% (w/v) agarose were derived from literature [10] and shown in Table S1. Hydraulic permeability parameters were *k*_0_ = 2.65 *×* 10^−12^ (m^4^/N s) and *M* = 4.1 .

#### S3.3. Boundary Conditions

The TE constructs were subject to mechanical loading in three different bioreactor setups that are commonly used in literature to apply unconfined compression, combined compression and shear, and hydrostatic pressure respectively. The boundary conditions applied to mimic mechanical loading for the first three cases were the same as those of the osteochondral plugs described in section 2.2 (boundary conditions **(a-b)**), with the only exception that the coefficient of friction between the compression plates (or rotating ball) and the TE construct was chosen to be 0.1 as derived from Geurds et al. [23].

##### (c) Hydrostatic Pressure

For knee articular cartilage *in vivo*, most of the compressive forces are transferred to hydrostatic pressure (HP) due to interstitial fluid pressurization, which acts as a primary load bearing mechanism in joints [24]. To harness this hydrostatic pressure to stimulate cartilage production in TE constructs, hydrostatic bioreactors have been used in literature imparting a wide range of loading magnitudes and frequencies [24, 25, 26]. In the present study, FE analysis of TE constructs subjected to static (0 Hz) and dynamic hydrostatic presure (at a frequency of 1 Hz) were performed. The loading magnitudes for hydrostatic pressure application were 0.5 MPa, 5 MPa, 10 MPa and 50 MPa, which were chosen to cover the wide range of magnitudes reported in literature [24]. The hydrostatic pressure was applied using a ‘Pressure Load’ in Abaqus.

#### S3.4. Implementation

The development and analysis of FE models were performed using Abaqus. To implement the fibril reinforced poroviscoelastic model of cartilage, multiple user-defined subroutines in Fortran were used. In order to run simulations in parallel across multiple cores, the UMAT subroutines were made thread-safe by removing common blocks and UEXTERNALDB subroutines were developed to implement collagen fibril orientations. These improvements allowed simulations to be run in parallel, significantly reducing computation time. All simulations were performed on the wICE high-performance computing (HPC) cluster at the Flemish Supercomputing Center (VSC), KU Leuven. The cluster consists of 172 IceLake nodes, each equipped with two Intel Xeon Platinum 8360Y CPUs (2.4 GHz, 36 cores per CPU, 1 NUMA domain, and 1 L3 cache per CPU), 256 GiB RAM, and 960 GB SSD local disk. The default memory per core is 3400 MiB. Job submissions were managed via Slurm scripts.

**Figure S4:**
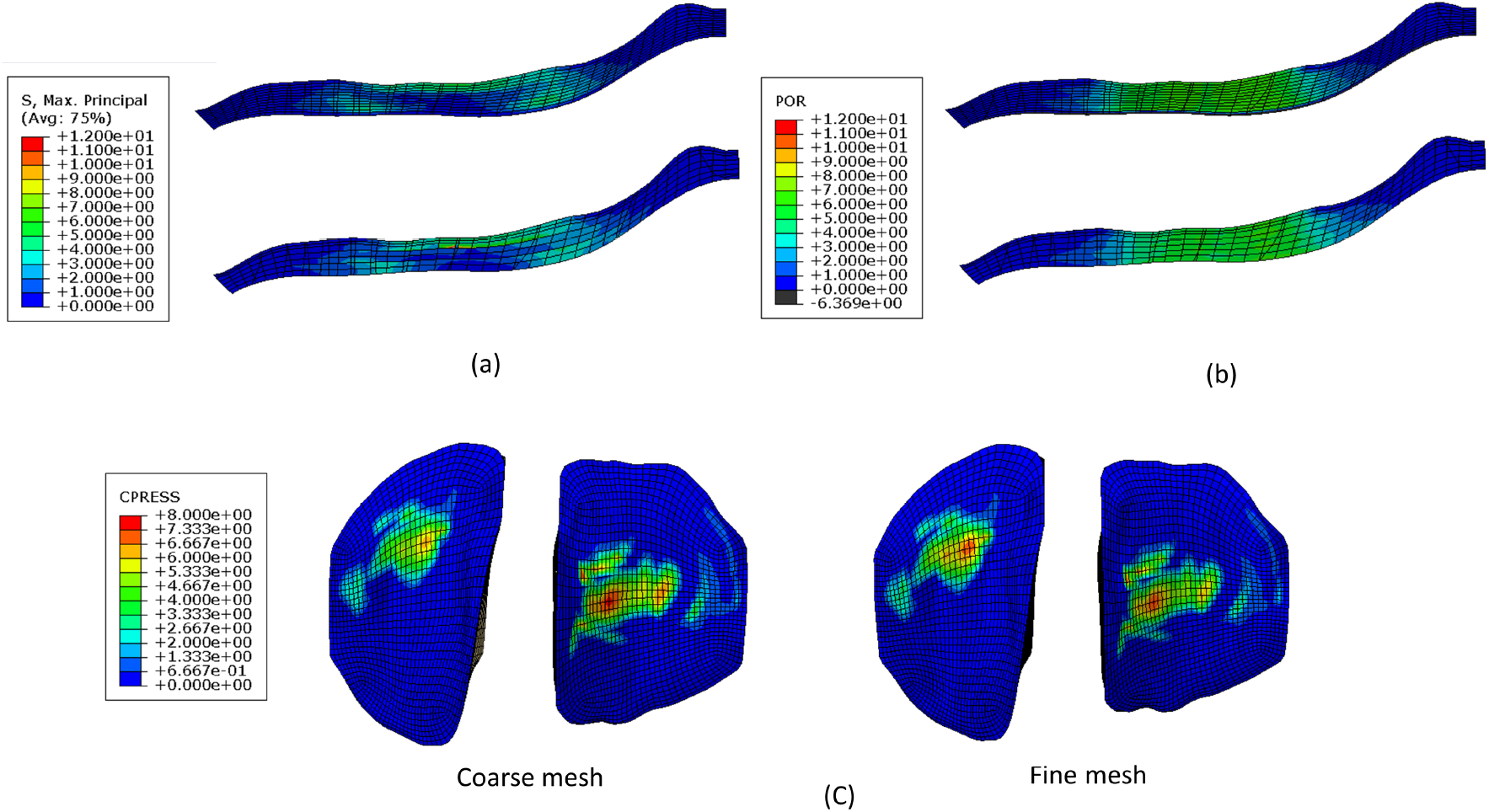
Mesh convergence study of knee joint. (a) Maximum principal stress. (b) Pore pressure. (c) Contact pressure

**Figure S5:**
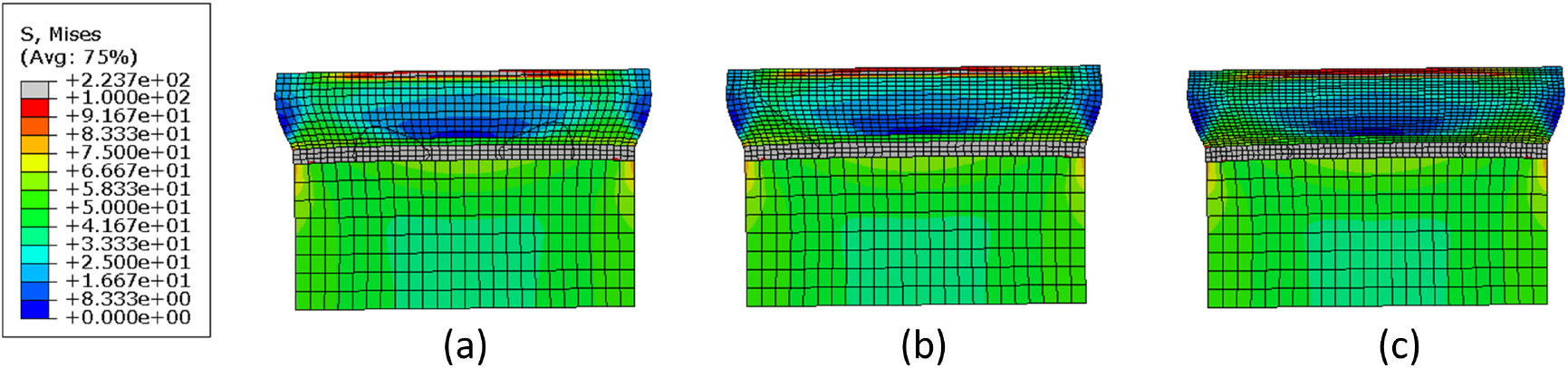
Mesh convergence study for osteochondral plugs at 30% strain. (a) Mesh=32023 elements. (b) Mesh=50263 elements. (c) Mesh=91447 elements.

**Figure S6:**
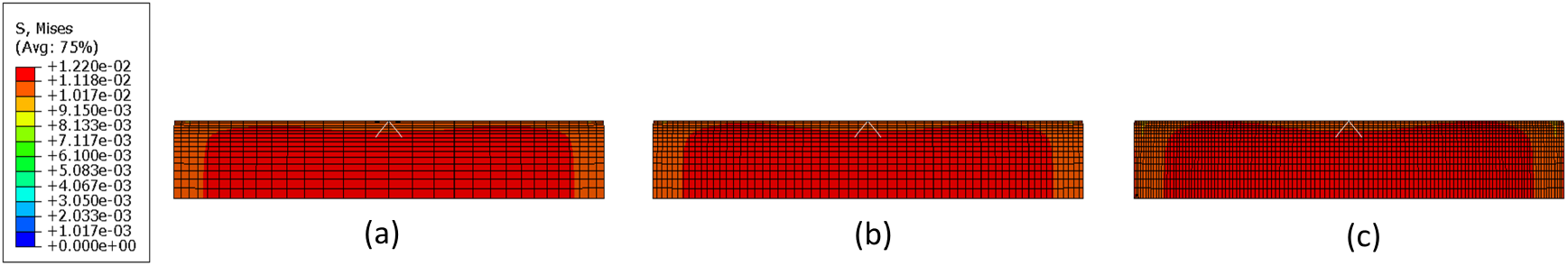
Mesh convergence study for agarose constructs 30% strain. (a) Mesh=18297 elements. Mesh=54672 elements. (c) Mesh=221127 elements.

**Figure S7:**
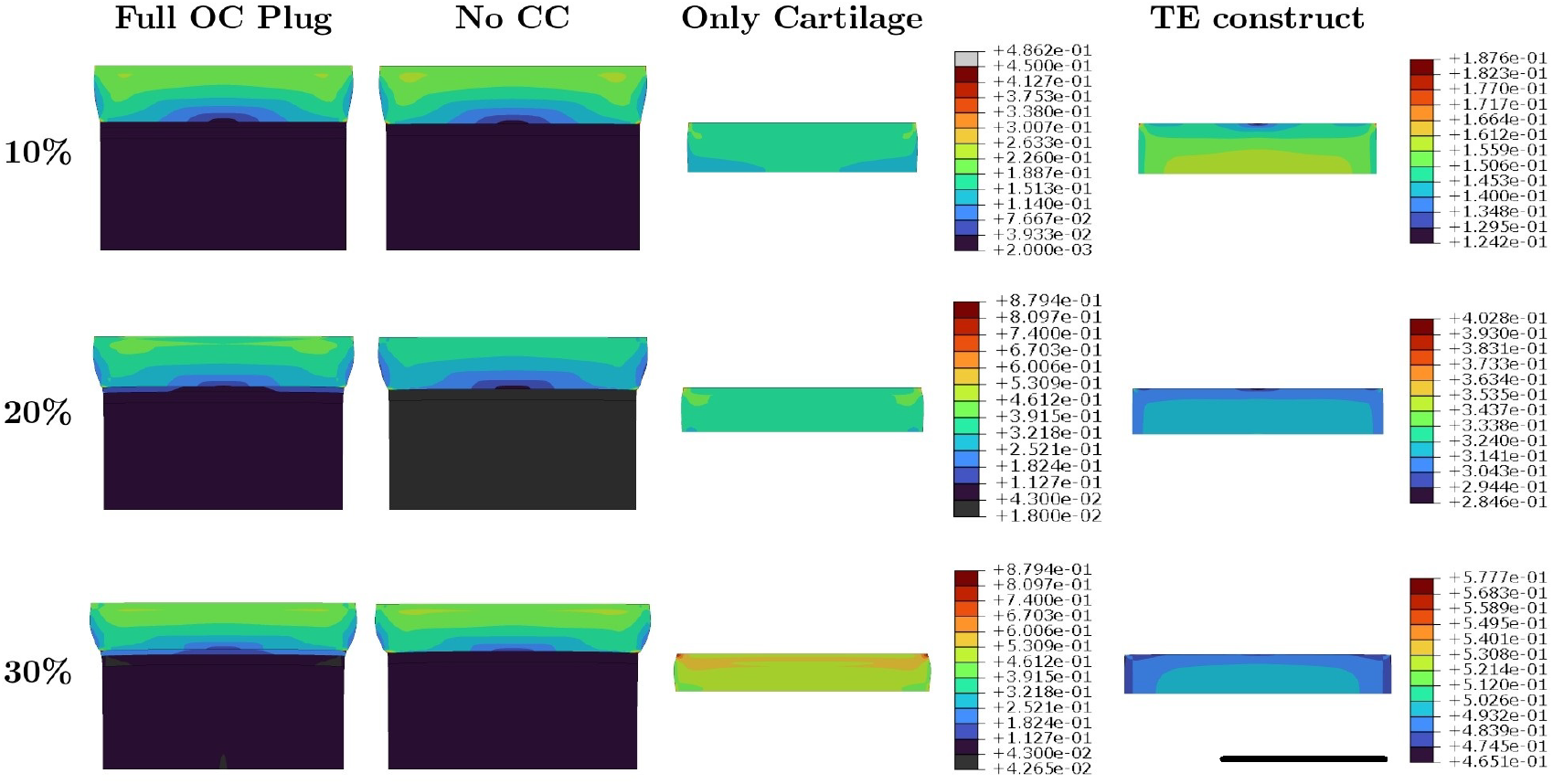
Distribution of maximum shear strain in different tissues of the OC plug and the TE construct for 10%, 20% and 30% strain applied in dynamic unconfined compression with 1Hz frequency, NoCC - No Calcified Cartilage, Scale bar = 5mm

**Figure S8:**
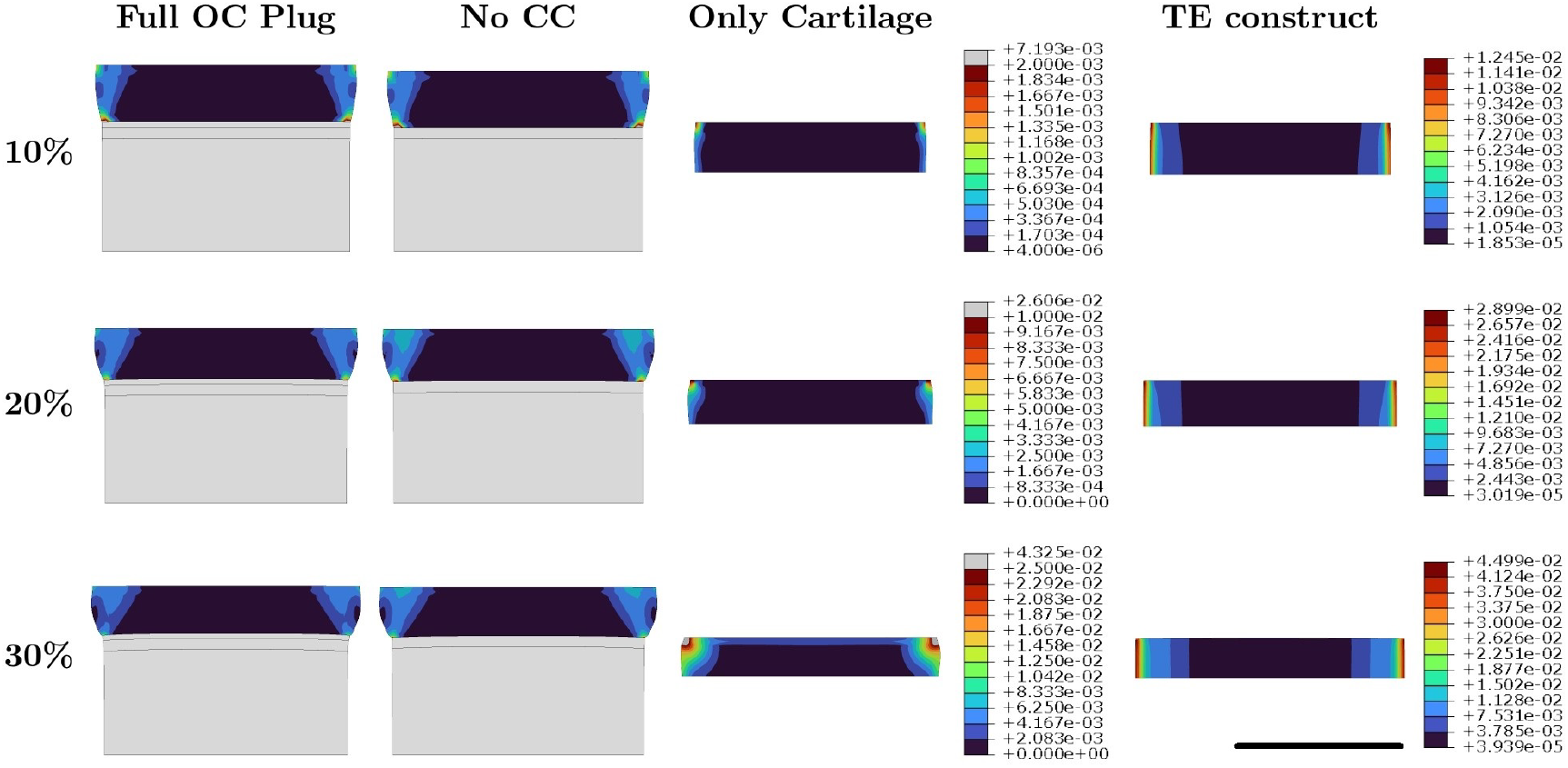
Distribution of pore fluid velocity (in mm/sec) in different tissues of the OC plug and the TE construct for 10%, 20% and 30% strain applied in dynamic unconfined compression with 1Hz frequency, NoCC - No Calcified Cartilage, Scale bar = 5mm

**Figure S9:**
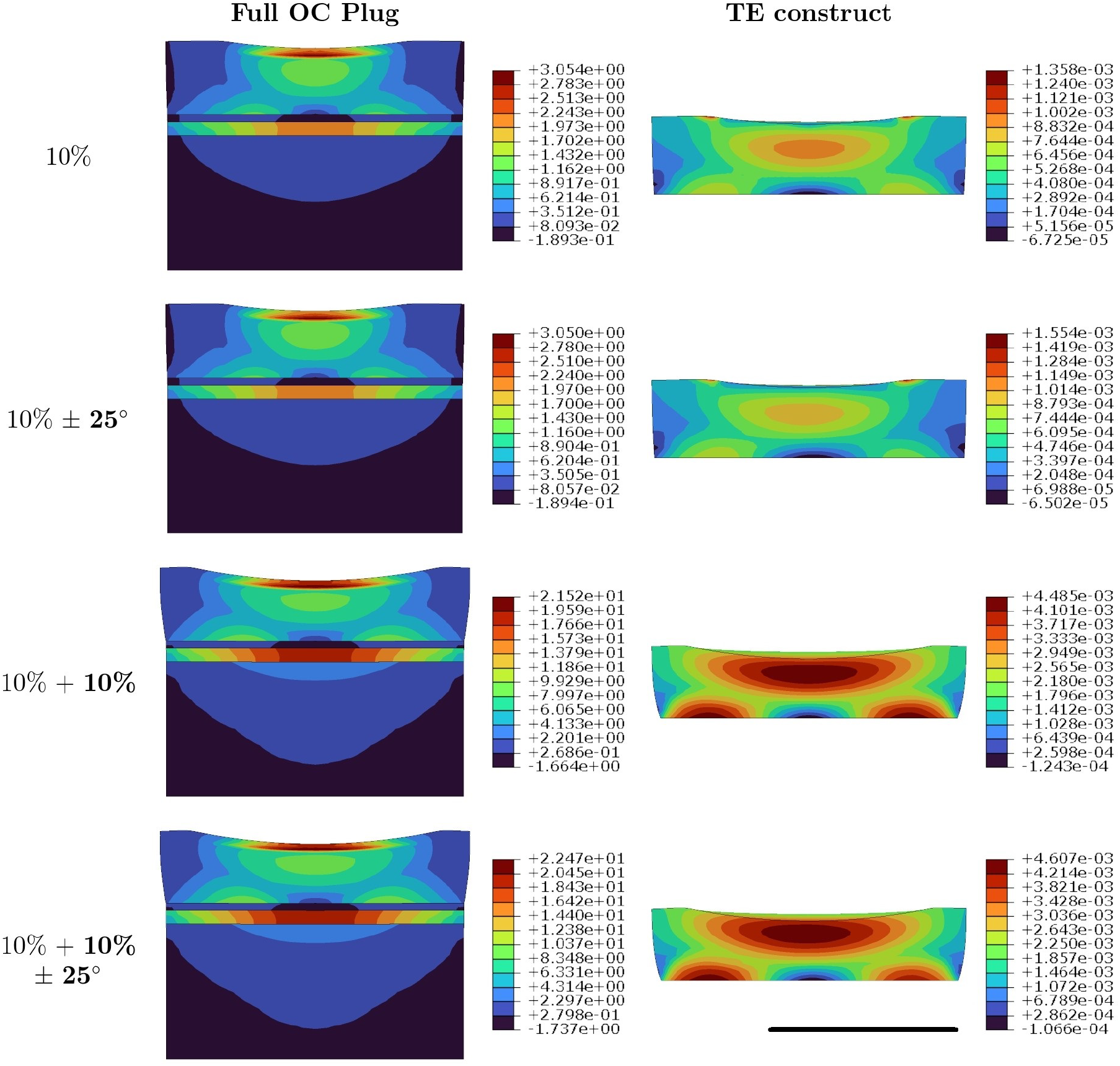
Distribution of maximum principal stress (in MPa) in OC plug and the TE construct for 4 different conditions of applied compression with ball rotation. Values in bold in the first column represents a dynamic boundary condition with 1Hz frequency, while normal fonts are for quasi-static. Scale bar = 5mm

**Figure S10:**
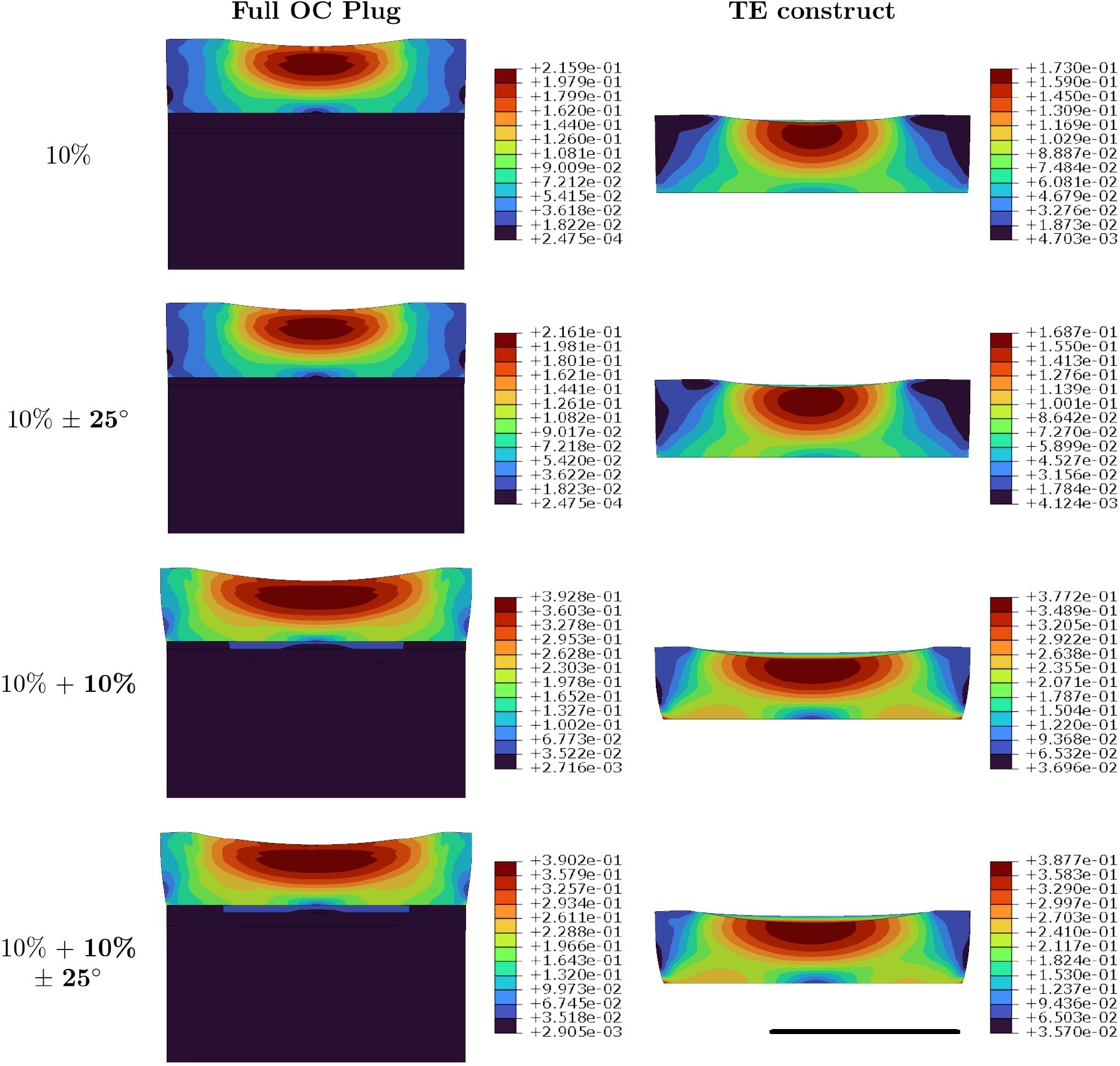
Distribution of maximum shear strain in OC plug and the TE construct for 4 different conditions of applied compression with ball rotation. Values in bold in the first column represents a dynamic boundary condition with 1Hz frequency, while normal fonts are for quasi-static. Scale bar = 5mm

**Figure S11:**
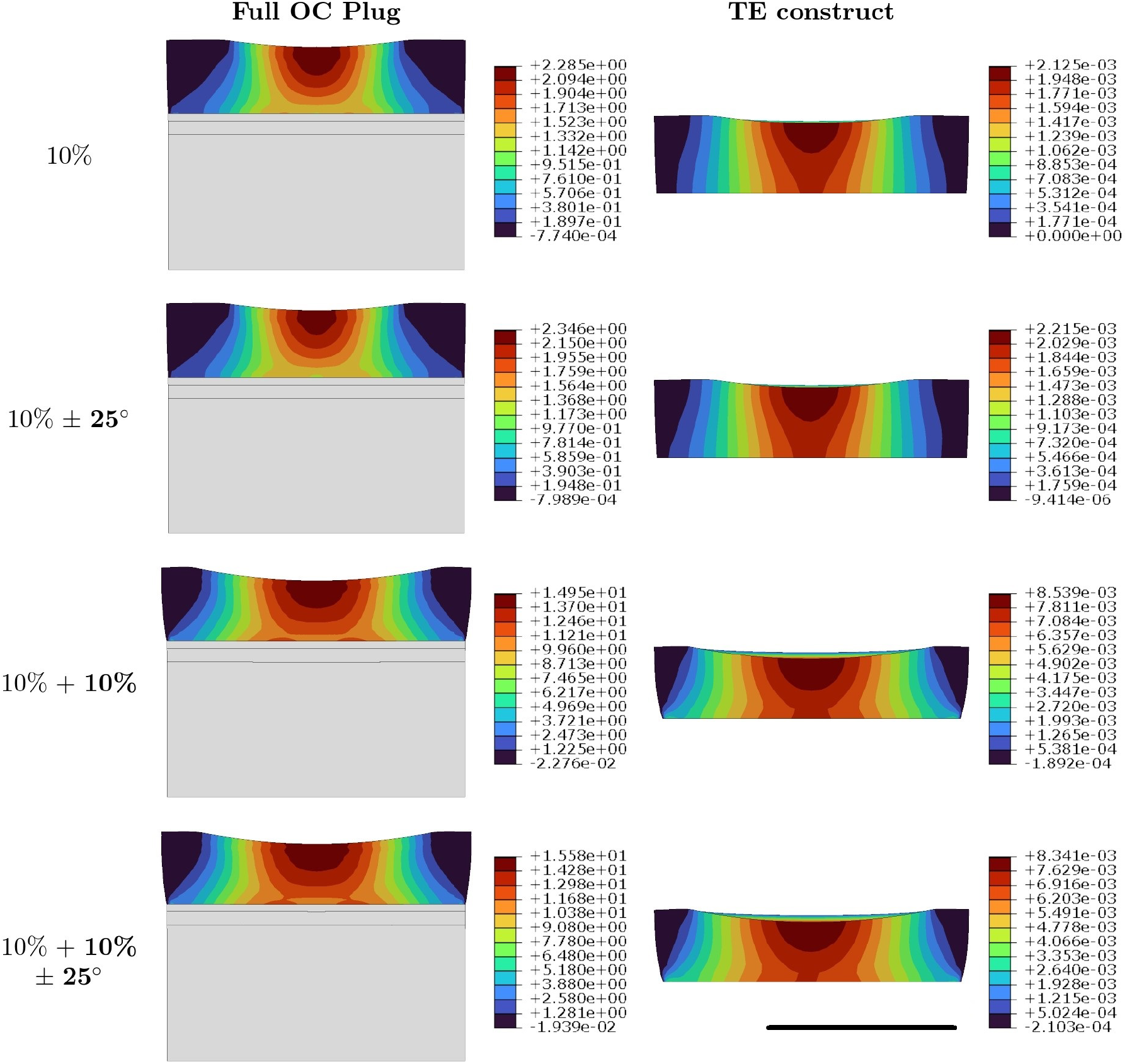
Distribution of pore fluid pressure (in MPa) in OC plug and the TE construct for 4 different conditions of applied compression with ball rotation. Values in bold in the first column represents a dynamic boundary condition with 1Hz frequency, while normal fonts are for quasi-static. Scale bar = 5mm

**Figure S12:**
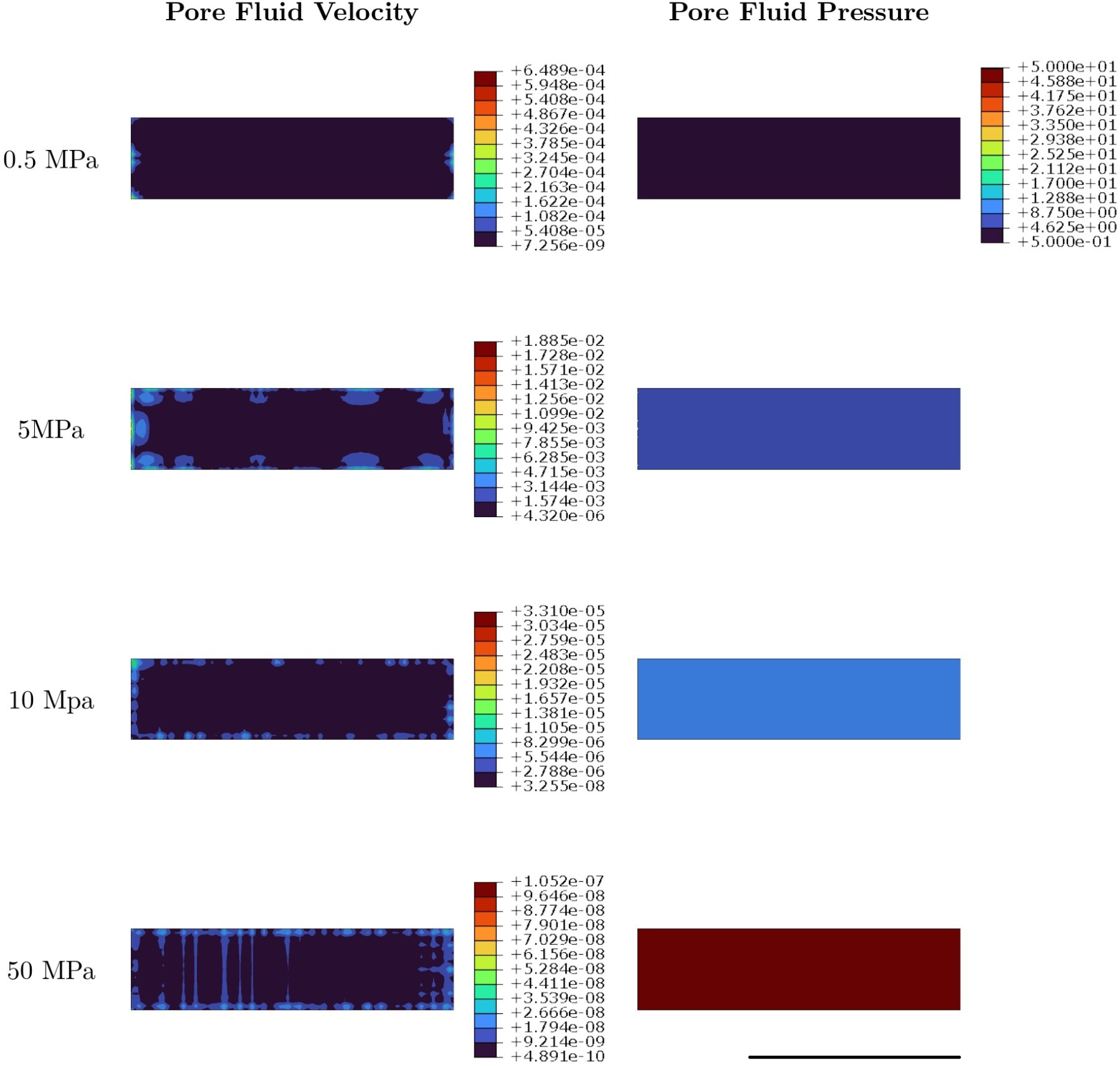
Distribution of pore fluid velocity (in mm/sec) and pore pressure (MPa)in TE construct for 4 different conditions of applied dynamic hydrostatic pressure at 1Hz frequency. Scale bar = 5mm

**Figure S13:**
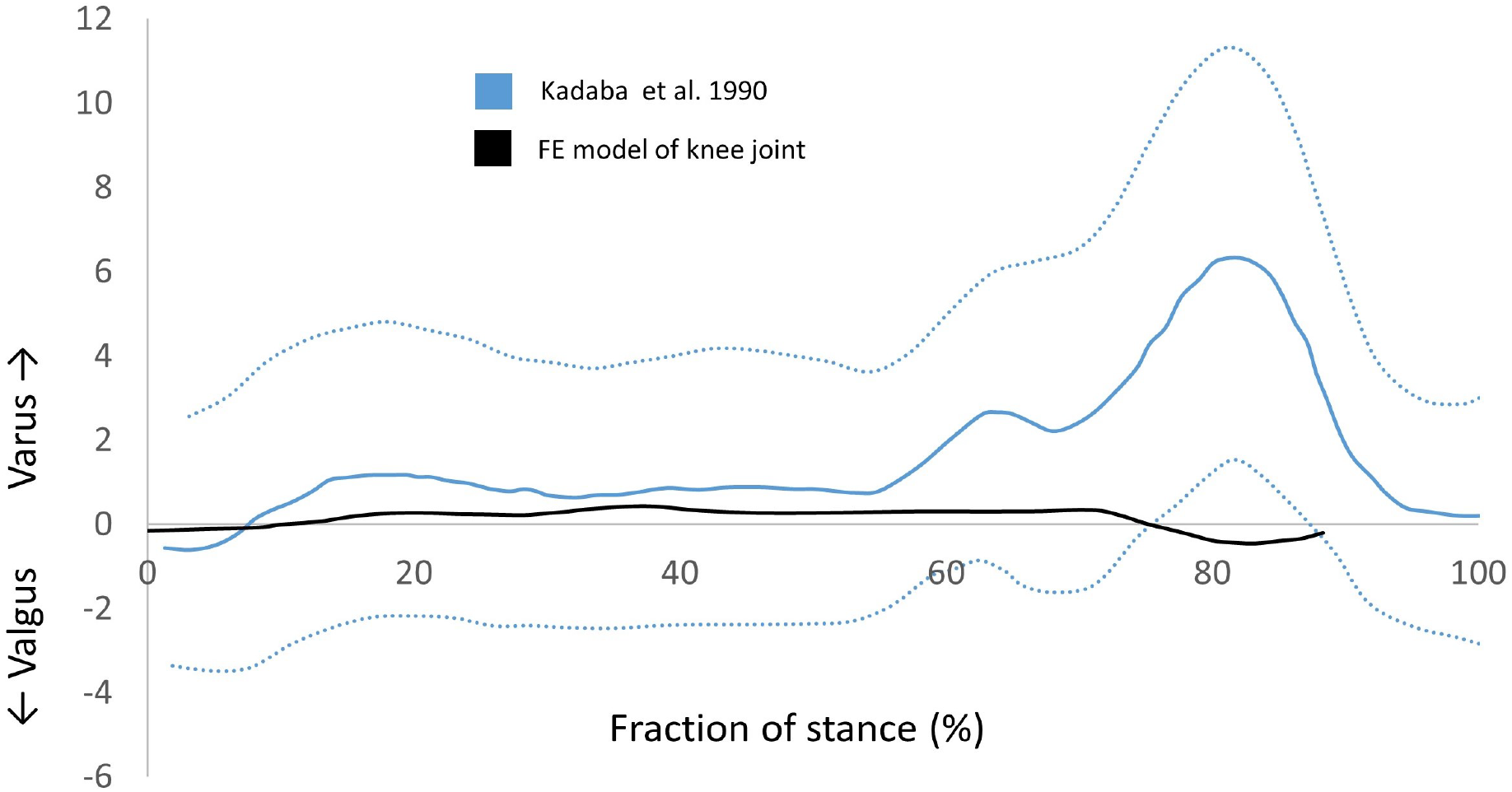
Comparison of varus-valgus rotation between the FE model and the experimental measurements as was done in Kadaba et al.[27]. The blue dotted lines represent the standard deviation

## Notes

### Competing Interest Statement

Wouter Wilson is an employee at Sioux Technologies, Netherlands

